# Mechanistically Interpretable Toxicity Prediction Through Multimodal Integration of Structure and Transcriptomics

**DOI:** 10.1101/2025.11.14.686754

**Authors:** Guillaume Cattebeke, Anne-Sofie Vermeersch, Davie Cappoen, Dieter Deforce, Filip Van Nieuwerburgh

## Abstract

Advances in computational toxicology increasingly emphasize the need for models that deliver both scalable predictive performance and mechanistic insights. Nevertheless, most approaches fall short of capturing the underlying mechanisms that drive toxicity. Herein, we describe a scalable multimodal modeling framework that integrates chemical fingerprints with high-throughput transcriptomic dose-response profiles across three human cell lines to predict activity for 41 curated Tox21 assay endpoints. Using gradient-boosted decision trees and nested compound-aware cross-validation, 13 assays achieved robust performance (mean AUPRC > 0.75), spanning nuclear receptor signaling, stress-response pathways, and xenobiotic metabolism. SHAP-based feature attribution analysis showed that predictions depend on both structural motifs and transcriptional programs, in a manner consistent with established mechanistic relationships between chemical structure, nuclear receptor biology, and adaptive cellular responses. These findings illustrate how structure-HTTr dose-response signature integration enables models that are accurate and mechanistically grounded, shifting computational toxicology toward transparent and biologically informed chemical risk evaluation.

## Introduction

The assessment of chemical toxicity and safety has traditionally relied heavily on animal testing. Yet, despite several significant contributions, animal models often remain poor predictors of human outcomes^1^. Translating their findings to human health has proven challenging, as inter-species differences in physiology, metabolism, and molecular signaling often result in poor concordance rates between animal test results and actual human responses. Moreover, traditional *in vivo* studies are increasingly criticized for their ethical implications, substantial costs, and poor reproducibility, limiting their relevance in modern-day toxicology^2^. At the same time, the expanding chemical landscape presents a different challenge, as humans are exposed to a wide range of compounds, characterized by distinct physicochemical properties, biological targets, and unique mechanisms of toxicity. Taken together, this creates a level of complexity that traditional testing methods struggle to address, making comprehensive evaluation both resource-intensive and often ineffective.

In light of this, alternative testing methods have progressed considerably^3,4^. Modern *in vitro* systems now enable the evaluation of large amounts of chemicals on biologically relevant mechanisms without relying on whole-animal models, and instead contribute to a growing framework of innovative strategies collectively named New Approach Methodologies (NAMs). Defined as any *in vitro, in silico*, or alternative testing method used to inform chemical hazard and risk assessment without the use of animals, NAMs have gained traction in recent years, with regulatory agencies beginning to incorporate them into established evaluation frameworks^5^. Within this context, high-throughput screening (HTS) platforms enable the evaluation of large and diverse sets of chemicals at an unprecedented scale. Likewise, high-throughput transcriptomics (HTTr) comprehensively profiles gene expression changes in response to a wide variety of chemical perturbations. As such, large-scale collaborative initiatives have begun to systematically generate, organize, and share high-quality toxicity data. A prominent example is the U.S. Environmental Protection Agency’s (EPA) Toxicity Forecaster (ToxCast) program, which compiles results from hundreds of high-throughput assays^6^. More recently, efforts at the EPA’s Center for Computational Toxicology and Exposure (CCTE) have focused on systematically cataloging chemical-induced gene expression changes through the adoption of a TempO-Seq–based targeted RNA-Seq platform, which enables cost-effective transcriptomic screening at scale^7–9^. Through large HTTr screenings, over a 1,000 chemicals from the ToxCast collection have now been profiled across multiple human cell lines, effectively expanding the chemical screening landscape with complementary transcriptome-wide gene expression data.

Although transcriptomic approaches have shown promise in specific case studies, their application to chemical toxicity assessment has generally been limited to specific biological domains or small chemical sets, often relying on reduced gene panels that capture only a subset of the transcriptional response^10,11^. In parallel, traditional computational toxicology has relied heavily on chemical descriptors and quantitative structure–activity relationship (QSAR) models, which capture intrinsic molecular features but lack direct biological context^12^. Although QSAR models can perform well within specific chemical domains, their reliance on structural descriptors limits mechanistic interpretability, highlighting the need for biological measurements that reflect the consequences of chemical exposure. Large-scale HTS and HTTr platforms now address this by enabling the systematic profiling of thousands of chemical–dose–cell line combinations, generating complementary information on both chemical activities and downstream cellular responses. Despite advances in modeling and rapid progress in generating these datasets, current approaches remain limited to narrow transcriptomic profiles, single-dose designs, or single-cell-line systems, leaving no general framework for integrating full transcriptome dose–response information with chemical structure at scale. Advancing beyond these limitations will require integrative approaches that extend biological coverage, uncover conserved mechanisms across diverse assays, and enable models that are both predictive and mechanistically informative.

In this study, we introduce a modeling framework that integrates these modalities and predicts bioactivity across Tox21 assay endpoints. The framework combines chemical fingerprints with transcriptomic dose– response summaries from multiple human cell lines to generate scalable, mechanistically interpretable predictions across a diverse panel of nuclear receptor, stress-response, and xenobiotic-metabolism assays. It demonstrates how large-scale data integration can enhance chemical safety assessment and shift predictive toxicology toward human-relevant, mechanism-based evaluation frameworks.

## Results

### ASSAY CURATION YIELDS 41 ENDPOINTS WITH COMPACT TRANSCRIPTOMIC FEATURE SPACE

To enable robust and generalizable predictive modeling, we first curated a high-confidence subset of endpoints from the Tox21 assay collection. Starting from 290 assay components, stepwise filtering based on sample size, class distribution, data completeness, and viability metrics retained 41 endpoints suitable for model development, collectively spanning key toxicological mechanisms such as nuclear receptor signaling (e.g., estrogen and androgen receptor activity), stress-response pathways (e.g., DNA damage and oxidative stress), and xenobiotic metabolism (e.g., cytochrome P450 inhibition) (Supplementary Table 1). Most exclusions resulted from initial data-quality filtering, which removed well over half of the original assay components as formats or readouts not suitable for primary activity modeling. Subsequent viability and class-balance criteria further narrowed the remaining set to those with sufficiently reliable and well-distributed activity data. Collectively, this curated panel offers coverage of toxicological mechanisms relevant to human chemical safety assessment.

The 41 endpoints cover 1,985 unique chemicals with complete HTTr measurements, corresponding to a fraction of the broader ToxCast chemical space across all Tox21 assays (8,948 unique DTXSIDs). While some endpoints exhibited nearly complete activity coverage, others displayed substantial data sparsity with an overall missingness ranging from 10% to 60%. This uneven coverage coincided with substantial variation in both the fraction of actives (class balance) and the absolute number of active compounds across assays (Figure 1). For example, CYP450 antagonist assays (e.g., P450 CYP2C9 antagonist assay) contained comparatively large positive classes, consistent with their broad substrate specificity and central role in xenobiotic metabolism^13^. In contrast, nuclear receptor and transcription factor assays (e.g., ERa antagonist assay) typically had fewer active compounds, which is expected given that only structurally specific molecules can modulate these pathways^14^. The number of active compounds varied across endpoints, corresponding to class balances between 3% and 60%. Such imbalance has important implications for predictive modeling, as highly imbalanced datasets can lead to models that disproportionately favor the majority (inactive) class, resulting in inflated accuracy and diminished sensitivity toward minority (active) compounds. These rare actives are often of greatest toxicological concern, as they reveal specific molecular interactions capable of disrupting key signaling or metabolic pathways, and therefore highlight why sensitivity, rather than accuracy, is the more meaningful performance metric in this context.

**Figure 1:**
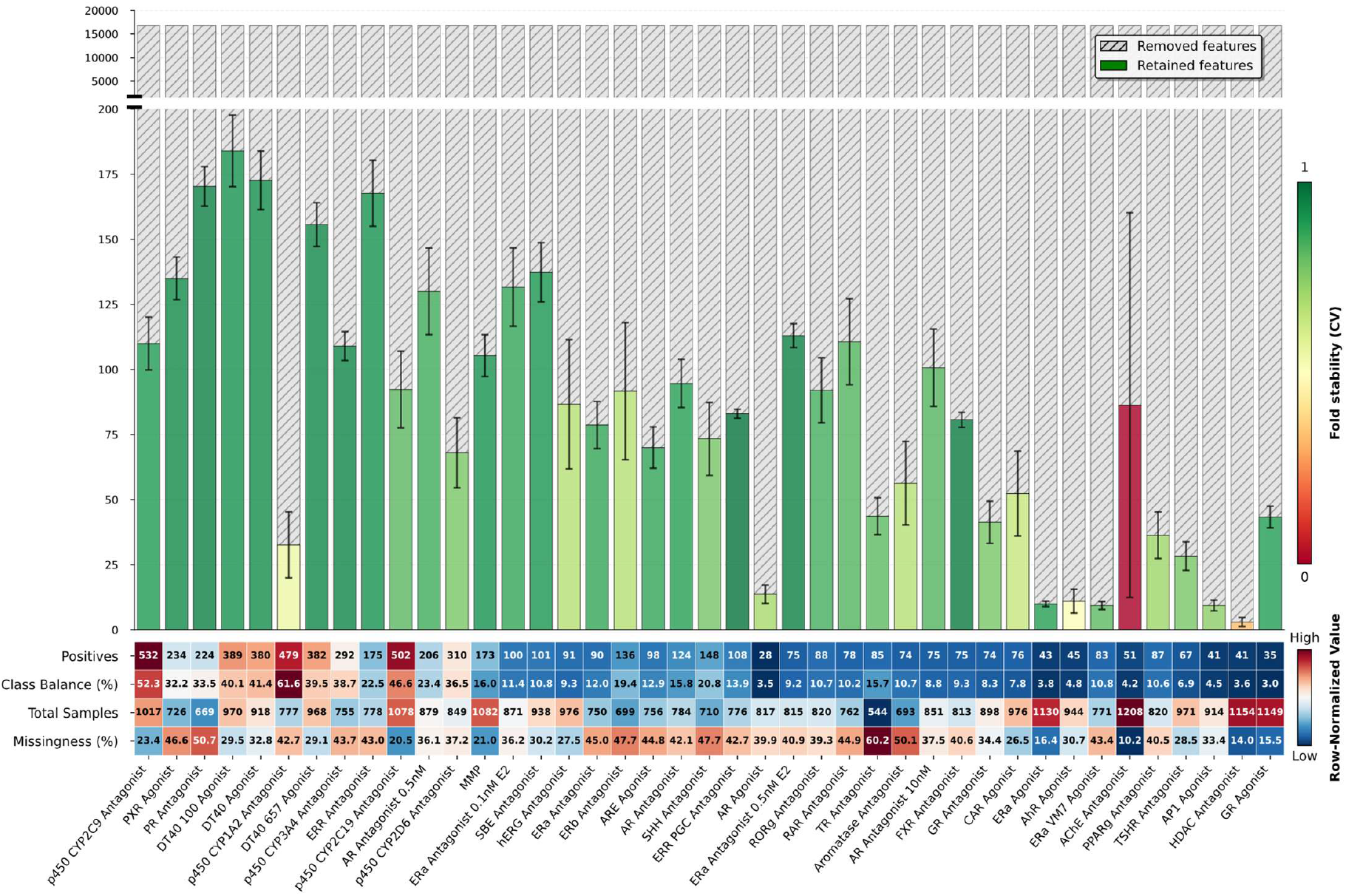
Feature reduction and dataset characteristics across 41 curated Tox21 assay endpoints. Bar heights represent the number of transcriptomic features retained after Boruta feature selection relative to the total feature space, and colors indicate cross-validation stability of retained features with greener bars denoting more consistent feature selection across folds. Error bars represent the standard deviation of feature counts across replicate Boruta runs. The lower heatmap summarizes mean dataset characteristics for each assay across 3 folds, including the number of positive samples, class balance, total sample size, and fraction of missing values. Each metric was row-normalized to emphasize within-metric variation across assays. Numerical annotations display the corresponding mean values.

To identify the transcriptomic features most informative for predicting bioactivity, we applied the Boruta feature-selection algorithm to the transcriptomic dose–response summary features for each endpoint. Boruta was chosen to address both the high dimensionality and the strong correlations inherent in these data. Unlike embedded or sparsity-based feature-selection approaches which typically yield a minimal subset of predictors optimized primarily for model performance, Boruta performs feature selection by retaining all predictors that demonstrate consistent importance, rather than restricting the model to the smallest set that maximizes predictive accuracy^15^. From an initial set of 16,821 dose-response summaries, the number of stably retained features varied across assays, with an average of 83 ± 50 per endpoint (Figure 1). This represents a reduction of more than two orders of magnitude in feature space, underscoring that only a small fraction of transcriptomic responses consistently carry predictive signal. Because the feature space consisted exclusively of pathway- and regulator-level summaries (originating from Hallmark, DoRothEA and PROGENy), the retained features reflected higher-order biological processes rather than individual gene responses. Retained features were distributed across these different summary layers, indicating that predictive signal arises from multiple pathways rather than a single dominant biological process. In parallel, chemical structure was encoded using the 166-bit MACCS fingerprints (Supplementary Table 2). Notably, the dimensionality of the Boruta-selected features was of the same order of magnitude as that of the MACCS fingerprints, which prevented one modality from dominating the feature space and facilitated more equal integration of chemical and biological descriptors during model training. In practice, this ensured that predictions were not biased toward structural similarity alone, but instead reflected both intrinsic chemical properties and observed cellular consequences. This balance is important, as prior multimodal profiling work has shown that structure- and biology-based descriptors tend to capture complementary signals and generally yield stronger and more mechanistically informative models when used together than when applied in isolation^16^. Consequently, the integrated feature space used here supports predictions shaped by both molecular scaffolds and the transcriptional programs they induce.

### AUPRC-FOCUSSED EVALUATION IDENTIFIES 13 PREDICTIVE ENDPOINTS

Model performance was evaluated on held-out test folds using a nested, group-aware cross-validation workflow to avoid information leakage. Overall, the integrated models achieved strong predictive performance across the 41 retained endpoints, although performance varied among assays (Table 1). Due to the pronounced class imbalance, metrics such as the area under the receiver operating characteristic curve (ROC AUC) can overestimate predictive ability, as high ROC AUC values may still correspond to poor sensitivity for the minority (active) class (e.g., GR Agonist assay). In contrast, the area under precision-recall curve (AUPRC) is more informative under class imbalance, as it explicitly quantifies the tradeoff between precision and recall for the positive class (i.e. active compounds). Accordingly, AUPRC was adopted as the primary metric for comparing model performance across endpoints. Sensitivity, specificity, and the Matthews correlation coefficient (MCC) were also evaluated to further characterize classifier behavior.

**Table 1:** Cross-validated predictive performance metrics of XGBoost models across 41 curated Tox21 assay endpoints. Results represent the mean ± standard deviation from 3-fold nested cross-validation. Metrics include ROC AUC (area under the receiver operating characteristic curve), AUPRC (area under the precision–recall curve), SE (sensitivity), SP (specificity), and MCC (Matthews correlation coefficient). Assays meeting the empirical predictability criterion (mean AUPRC > 0.75) are indicated with an asterisk and were selected for subsequent analyses.

| AEID | ASSAY | ROC AUC | AUPRC | SE | SP | MCC |
| --- | --- | --- | --- | --- | --- | --- |
| 3187 | p450 CYP2C9 Antagonist* | 0.896 ± 0.019 | 0.908 ± 0.016 | 0.846 ± 0.025 | 0.829 ± 0.044 | 0.645 ± 0.042 |
| 2363 | PXR Agonist* | 0.944 ± 0.016 | 0.899 ± 0.020 | 0.827 ± 0.009 | 0.841 ± 0.017 | 0.741 ± 0.015 |
| 2127 | PR Antagonist* | 0.939 ± 0.006 | 0.892 ± 0.014 | 0.866 ± 0.023 | 0.804 ± 0.030 | 0.738 ± 0.021 |
| 2130 | DT40 100 Agonist* | 0.911 ± 0.021 | 0.889 ± 0.025 | 0.838 ± 0.035 | 0.822 ± 0.015 | 0.710 ± 0.027 |
| 1134 | DT40 Agonist* | 0.902 ± 0.024 | 0.886 ± 0.030 | 0.792 ± 0.051 | 0.839 ± 0.018 | 0.682 ± 0.047 |
| 3188 | p450 CYP1A2 Antagonist* | 0.827 ± 0.017 | 0.883 ± 0.011 | 0.877 ± 0.069 | 0.768 ± 0.062 | 0.454 ± 0.044 |
| 2131 | DT40 657 Agonist* | 0.891 ± 0.013 | 0.872 ± 0.013 | 0.805 ± 0.027 | 0.775 ± 0.030 | 0.633 ± 0.020 |
| 2544 | p450 CYP3A4 Antagonist* | 0.893 ± 0.018 | 0.849 ± 0.025 | 0.802 ± 0.027 | 0.772 ± 0.041 | 0.643 ± 0.027 |
| 2057 | ERR Antagonist* | 0.935 ± 0.026 | 0.835 ± 0.039 | 0.795 ± 0.080 | 0.749 ± 0.032 | 0.698 ± 0.064 |
| 3186 | p450 CYP2C19 Antagonist* | 0.843 ± 0.029 | 0.834 ± 0.018 | 0.782 ± 0.021 | 0.771 ± 0.048 | 0.582 ± 0.034 |
| 1816 | AR Antagonist 0.5nM* | 0.917 ± 0.015 | 0.795 ± 0.038 | 0.736 ± 0.045 | 0.747 ± 0.020 | 0.658 ± 0.028 |
| 3185 | p450 CYP2D6 Antagonist* | 0.842 ± 0.022 | 0.794 ± 0.028 | 0.753 ± 0.045 | 0.687 ± 0.053 | 0.542 ± 0.052 |
| 1854 | MMP* | 0.929 ± 0.008 | 0.767 ± 0.009 | 0.691 ± 0.038 | 0.743 ± 0.030 | 0.648 ± 0.014 |
| 2053 | ERa Antagonist 0.1nM E2 | 0.935 ± 0.008 | 0.720 ± 0.055 | 0.774 ± 0.016 | 0.619 ± 0.078 | 0.620 ± 0.022 |
| 2372 | SBE Antagonist | 0.916 ± 0.014 | 0.718 ± 0.058 | 0.652 ± 0.101 | 0.736 ± 0.116 | 0.636 ± 0.075 |
| 3210 | hERG Antagonist | 0.932 ± 0.028 | 0.705 ± 0.045 | 0.753 ± 0.080 | 0.582 ± 0.049 | 0.607 ± 0.055 |
| 786 | ERa Antagonist | 0.926 ± 0.010 | 0.684 ± 0.048 | 0.684 ± 0.061 | 0.657 ± 0.075 | 0.621 ± 0.031 |
| 2119 | ERb Antagonist | 0.893 ± 0.032 | 0.683 ± 0.095 | 0.752 ± 0.093 | 0.628 ± 0.081 | 0.599 ± 0.095 |
| 1110 | ARE Agonist | 0.904 ± 0.018 | 0.682 ± 0.045 | 0.670 ± 0.112 | 0.629 ± 0.029 | 0.562 ± 0.026 |
| 762 | AR Antagonist | 0.917 ± 0.003 | 0.677 ± 0.039 | 0.784 ± 0.074 | 0.582 ± 0.054 | 0.592 ± 0.013 |
| 2108 | SHH Antagonist | 0.859 ± 0.019 | 0.675 ± 0.043 | 0.670 ± 0.071 | 0.637 ± 0.032 | 0.537 ± 0.051 |
| 2070 | ERR PGC Antagonist | 0.866 ± 0.022 | 0.630 ± 0.026 | 0.641 ± 0.061 | 0.587 ± 0.063 | 0.543 ± 0.043 |
| 761 | AR Agonist | 0.902 ± 0.023 | 0.619 ± 0.041 | 0.529 ± 0.017 | 0.841 ± 0.059 | 0.600 ± 0.027 |
| 789 | ERa Antagonist 0.5nM E2 | 0.921 ± 0.004 | 0.616 ± 0.068 | 0.683 ± 0.116 | 0.628 ± 0.123 | 0.569 ± 0.061 |
| 1661 | RORg Antagonist | 0.890 ± 0.000 | 0.610 ± 0.114 | 0.673 ± 0.101 | 0.571 ± 0.129 | 0.547 ± 0.061 |
| 1839 | RAR Antagonist | 0.921 ± 0.025 | 0.609 ± 0.039 | 0.699 ± 0.056 | 0.614 ± 0.053 | 0.597 ± 0.034 |
| 804 | TR Antagonist | 0.867 ± 0.022 | 0.609 ± 0.027 | 0.633 ± 0.108 | 0.571 ± 0.051 | 0.500 ± 0.021 |
| 767 | Aromatase Antagonist | 0.900 ± 0.045 | 0.606 ± 0.099 | 0.614 ± 0.021 | 0.565 ± 0.078 | 0.517 ± 0.039 |
| 765 | AR Antagonist 10nM | 0.910 ± 0.014 | 0.589 ± 0.093 | 0.675 ± 0.061 | 0.554 ± 0.153 | 0.546 ± 0.057 |
| **1120** | *FXR Antagonist* | 0.912 ± 0.006 | 0.587 ± 0.045 | 0.650 ± 0.171 | 0.562 ± 0.079 | 0.523 ± 0.035 |
| **794** | *GR Antagonist* | 0.906 ± 0.019 | 0.569 ± 0.088 | 0.672 ± 0.087 | 0.478 ± 0.037 | 0.516 ± 0.058 |
| **2047** | *CAR Agonist* | 0.924 ± 0.027 | 0.545 ± 0.038 | 0.657 ± 0.066 | 0.565 ± 0.023 | 0.557 ± 0.042 |
| **785** | *ERα Agonist* | 0.884 ± 0.053 | 0.533 ± 0.129 | 0.516 ± 0.138 | 0.598 ± 0.117 | 0.513 ± 0.101 |
| **806** | *AhR Agonist* | 0.901 ± 0.033 | 0.521 ± 0.048 | 0.491 ± 0.025 | 0.616 ± 0.032 | 0.509 ± 0.003 |
| **788** | *ERα VM7 Agonist* | 0.825 ± 0.008 | 0.491 ± 0.018 | 0.540 ± 0.093 | 0.527 ± 0.016 | 0.461 ± 0.048 |
| **2210** | *AChE Antagonist* | 0.885 ± 0.033 | 0.453 ± 0.089 | 0.468 ± 0.172 | 0.594 ± 0.137 | 0.419 ± 0.110 |
| **1127** | *PPARγ Antagonist* | 0.859 ± 0.021 | 0.407 ± 0.039 | 0.552 ± 0.096 | 0.406 ± 0.020 | 0.385 ± 0.053 |
| **2040** | *TSHR Antagonist* | 0.817 ± 0.031 | 0.376 ± 0.062 | 0.371 ± 0.050 | 0.445 ± 0.092 | 0.351 ± 0.022 |
| **1846** | *AP1 Agonist* | 0.874 ± 0.040 | 0.367 ± 0.089 | 0.402 ± 0.101 | 0.525 ± 0.118 | 0.360 ± 0.046 |
| **2061** | *HDAC Antagonist* | 0.864 ± 0.025 | 0.326 ± 0.135 | 0.319 ± 0.103 | 0.529 ± 0.218 | 0.313 ± 0.103 |
| **793** | *GR Agonist* | 0.880 ± 0.025 | 0.304 ± 0.062 | 0.499 ± 0.080 | 0.379 ± 0.076 | 0.363 ± 0.098 |

The mean AUPRC values per assay ranged from near-random (AUPRC ≈ 0.3-0.5) for the most challenging nuclear receptor endpoints, to very high accuracies for the best-performing assays (AUPRC ≈ 0.8-0.9). As expected, predictive performance was strongly associated with the number of positive classes (i.e. active compounds) (Figure 2). This relationship is also reflected in the threshold-dependent sensitivity across training folds, as endpoints with higher AUPRC values typically achieved sensitivities above 0.8 at their F1-optimal thresholds. Consistent with this pattern, all of the CYP450 antagonism endpoints ranked among the most accurately predicted and had comparatively large numbers of positive classes. For instance, the CYP2C9 antagonist assay achieved an AUROC of 0.896 ± 0.019 and an AUPRC of 0.908 ± 0.016, and had the highest number of active compounds (n = 1,596), while assays such as the GR agonist (n = 105) and HDAC antagonist assay (n = 124) with one of the fewest number of active compounds, exhibited substantially lower performance.

**Figure 2:**
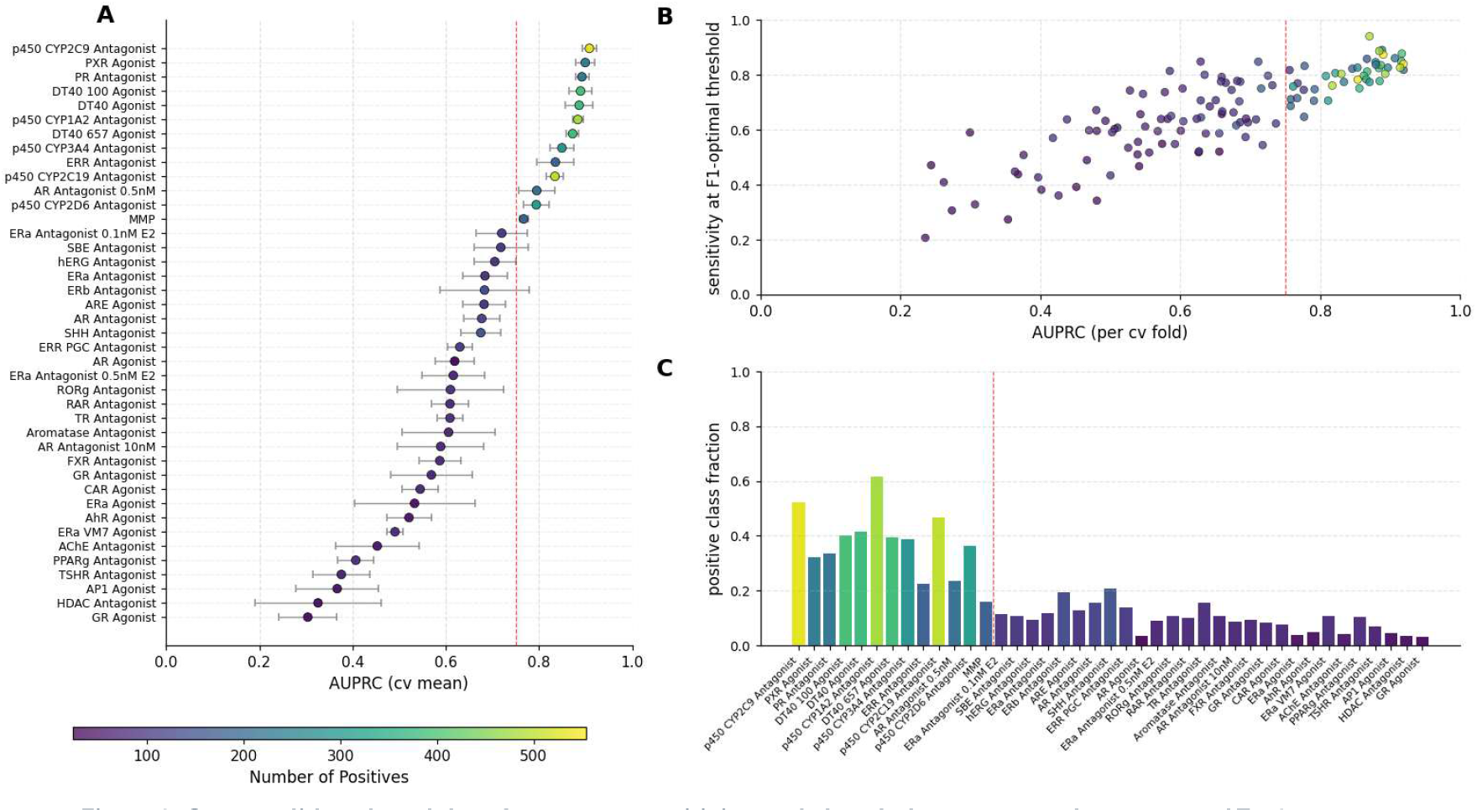
Cross-validated model performance, sensitivity, and class balance across the 41 curated Tox21 assays. **(A)** Each point shows the mean area under the precision–recall curve (AUPRC) for individual assays, with horizontal error bars indicating the standard deviation across cross-validation folds. Point colors reflect the mean number of positive samples in the test sets, illustrating variation in class balance across assays. The vertical dashed red line (AUPRC = 0.75) denotes the empirical performance threshold used to identify well-performing models for downstream analyses. **(B)** Relationship between fold-level AUPRC and sensitivity at the F_1_-optimal decision threshold. Each point represents a single cross-validation fold, colored by the number of positive samples in that fold. The vertical dashed red line marks the same AUPRC = 0.75 reference threshold. **(C)** Fraction of positive samples per assay, colored according to the shared “number of positives” color scale. Assays are sorted based on mean AUPRC values.

To delineate which endpoints can be considered reliably predictable, we applied an empirical cutoff of mean AUPRC > 0.75. This threshold reflects a clear separation from random performance and denotes models with a practically meaningful precision–recall trade-off (high recall at manageable false-positive rates) for identifying active compounds. From a toxicological perspective, emphasizing AUPRC is particularly appropriate, since positive (active/toxic) compounds are typically rare yet carry the greatest regulatory and health relevance. Using this criterion, 13 of the 41 endpoints were classified as predictive, representing assays with sufficiently balanced activity and mechanistic signal to support robust, generalizable insights.

### SHAP-BASED MODEL INTERPRETATION HIGHLIGHTS STRUCTURAL AND TRANSCRIPTOMIC DETERMINANTS OF BIOLOGICAL ACTIVITY

To identify which features most strongly contributed to model predictions, SHAP (SHapley Additive exPlanations) values were computed on the held-out test set within each cross-validation fold and were aggregated across folds. The resulting feature importance patterns indicated that model predictions were not driven by structural information alone, as the top contributors repeatedly combined MACCS fragments with dose-response summary features (Figure 3). Within the 13 selected assays, consistent patterns emerged (Supplementary Fig. 1-13). On the structural side, recurrent MACCS keys highlighted common molecular scaffolds and substituents across endpoints. The most influential fragments included six-membered ring motifs and oxygen-rich substituents, such as hydroxyl, ether-like, and other polar oxygen-containing groups (MACCS_145, MACCS_146, MACCS_140, MACCS_159, MACCS_126, MACCS_113), as well as amine- and heteroatom-containing features (MACCS_151, MACCS_124), and exhibited largely consistent predictive contributions. Together, these patterns are consistent with established principles of ligand–protein recognition: ring scaffolds often provide hydrophobic anchoring elements within binding pockets, while polar substituents enable hydrogen bonding and modulate the electronic properties of the molecular framework, enhancing both affinity and selectivity. Likewise, heteroatoms and amine functionalities introduce polarity and electrostatic complementarity that stabilize interactions within protein binding sites^17^. Beyond structural determinants, SHAP analysis also highlighted contributions from transcriptomic features derived from dose–response profiles. These features frequently captured conserved programs of stress adaptation and growth regulation, including hypoxia and ATF4-mediated integrated stress responses, IGF/ERK-driven survival signaling, and amino acid–sensing pathways. The recurrence of these modules across most assays suggests that the models capture central stress- and growth-regulatory circuits engaged during xenobiotic exposure^18,19^. At the same time, these signatures may also reflect more general cellular stress responses to xenobiotic exposure, and therefore may not always correspond to assay-specific mechanisms. Nonetheless, several assays showed transcriptomic signatures that were unique to their specific biological mechanisms.

**Figure 3:**
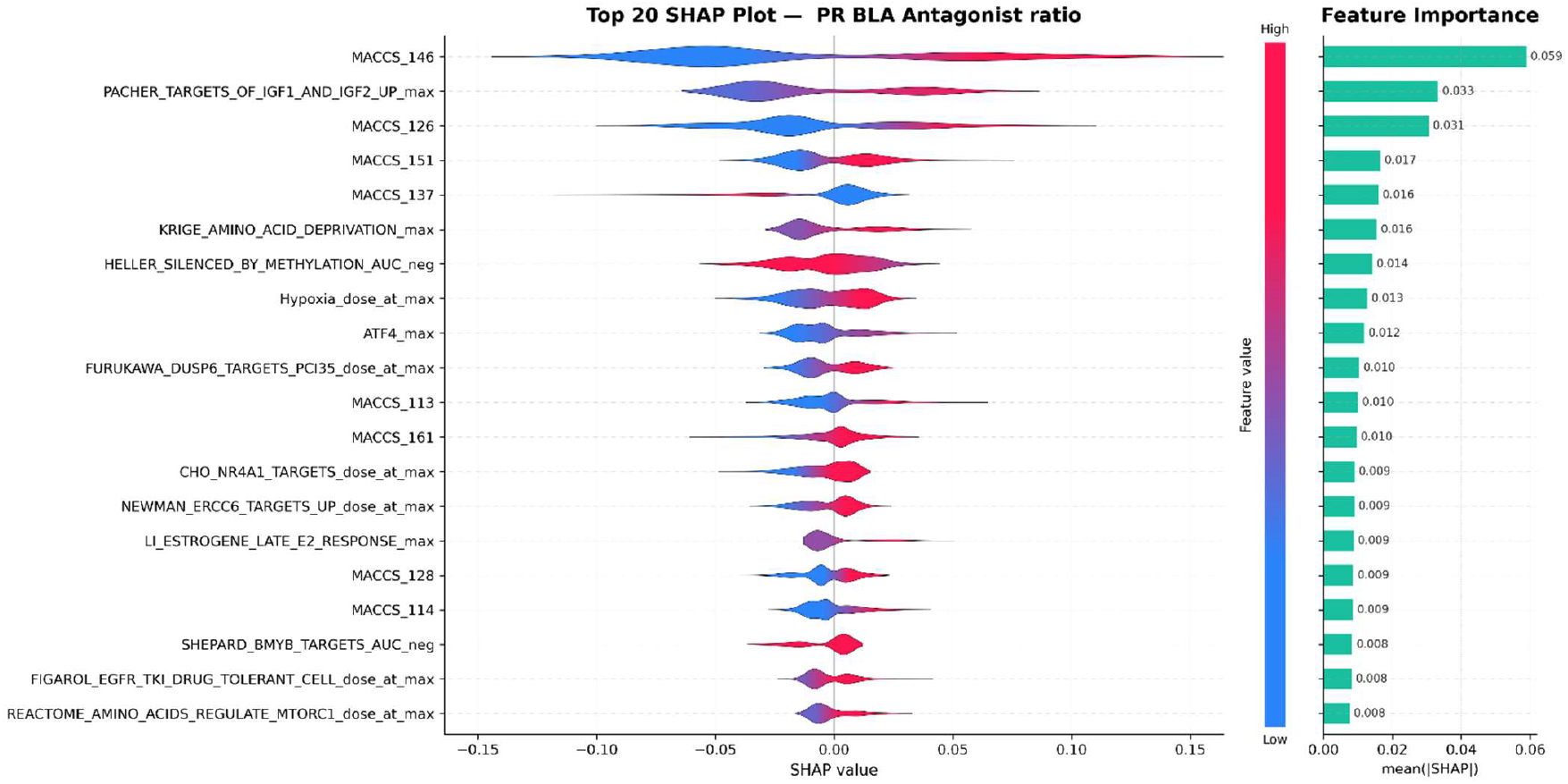
SHAP analysis reveals key molecular and biological features driving progesterone receptor (PR) beta-lactamase (BLA) antagonist activity prediction. The violin plots (left) display the distribution of SHAP values for the top 20 most important features, ordered by mean absolute SHAP value (right). Each violin’s width represents the density of SHAP values across all samples, with the x-axis indicating feature contribution magnitude and direction. The color gradient within violins encodes the normalized feature value revealing feature-prediction relationships. The feature set encompasses structural descriptors (MACCS fingerprints), transcriptional signatures (e.g., Hallmark pathways) and pathway activities (e.g., DoRothEA regulons). Feature suffixes indicate aggregation methods: _max (maximum value), _dose_at_max (dose at maximum response), and _AUC_neg (area under curve for negative enrichment, with flipped sign for interpretation).

To illustrate these trends, the progesterone receptor (PR) beta-lactamase (BLA) antagonist assay model was selected as a representative case for SHAP-based interpretation (Figure 3). Although not the top-performing assay based on AUPRC, it highlights a well-characterized nuclear receptor pathway for which structural determinants and transcriptomic profiles are well understood, making it easier to contextualize SHAP values in terms of mechanistic relevance and biological interpretability. Within this model, several structural features stood out as consistent with known pharmacology of antiprogestins. For instance, MACCS keys capturing oxygen-rich and heteroatom-containing fragments, including multiple oxygen atoms and ether-like linkages (MACCS_146, MACCS_126, MACCS_113) and amine or ring-nitrogen features (MACCS_151, MACCS_161, MACCS_137), were positive contributors toward antagonism, likely reflecting the 11β-aryl and heteroatom substitutions that distinguish antagonists such as mifepristone from receptor agonists^20^. SHAP analysis also identified transcriptomic programs closely linked to PR biology. The most direct PR-relevant feature was the late estrogen response module (LI_ESTROGENE_LATE_E2_RESPONSE_max), which contributed positively towards antagonism, a result that likely reflects compensatory activation of estrogen-associated pathways through ER–PR crosstalk^21^. Additional features captured compensatory growth factor signaling, most notably the upregulation of IGF1/2 targets (PACHER_TARGETS_OF_IGF1_AND_IGF2_UP_max) and modulation of ERK-related programs (FURUKAWA_DUSP6_TARGETS_PC3S_dose_at_max). Compounds that activated ERK/DUSP6 modules only at higher concentrations were more likely to be classified as PR antagonists, suggesting that MAPK engagement represents a secondary adaptative response rather than a direct effect. This interpretation aligns with mechanistic studies showing that while PR antagonists act directly at the receptor, ERK/DUSP6 feedback emerges later as a compensatory mechanism to sustain cell survival^22^. Collectively, the PR model exemplifies how structural motifs, generic stress-adaptation pathways, and receptor-specific transcriptomic signatures converge to produce mechanistically interpretable predictions that align with established principles of receptor pharmacology.

## Discussion

Integrating high-throughput transcriptomic data with large-scale chemical screening provides a path toward predictive toxicology that is both mechanistically informed and scalable across diverse chemical spaces. Despite major advances in high-throughput screening (HTS) technologies, most existing assays remain narrowly focused on predefined molecular targets or signaling pathways, offering only a partial view of cellular toxicity mechanisms. This target-centric design has produced extensive yet fragmented datasets that capture activity against specific pathways, but overlook the broader transcriptional and adaptive responses that determine cellular fate. As a result, computational models predicting HTS data often depend primarily on chemical structure information, which lacks direct connection to the biological mechanisms that underlie toxicity. Conventional quantitative structure–activity relationship (QSAR) approaches therefore remain limited in their ability to explain how molecular features translate into cellular outcomes. High-throughput transcriptomics (HTTr) addresses this gap by capturing transcriptome-wide molecular consequences of chemical exposure, revealing both direct target engagement and downstream adaptive responses. When integrated with structural information, transcriptomic features enable models to link molecular properties with the biological programs they trigger, transforming predictive toxicology from a structure-based inference exercise into a mechanistically grounded modeling framework. In this study, we extend this concept by integrating chemical fingerprints with full transcriptomic dose–response profiles across three human cell lines. This combination provides a level of biological breadth and mechanistic resolution not achieved in prior HTTr–QSAR studies, and mitigates the limitations of either modality alone, consistent with large-scale toxicogenomic studies where similar strategies improved toxicity prediction and interpretability^16,23,24^.

A persistent challenge in HTS data interpretation lies in distinguishing specific biological activity from nonspecific cytotoxicity. Many Tox21 assays are sensitive to stress-induced signaling, leading to widespread activation across multiple endpoints once concentrations approach cytotoxic thresholds, a phenomenon commonly referred to as the cytotoxicity burst^6,25^. In this sense, cytotoxicity is both an important biological endpoint and a potential confounding factor for mechanistic interpretation. In some cases, the majority of apparent positives can be attributed to this effect rather than true target modulation^26^. These cytotoxicity-driven activations introduce systematic noise that complicates the separation of specific mechanisms from global stress responses. High-throughput transcriptomics provides a way to disentangle these effects: gene expression profiles capture both general stress signatures and pathway-specific perturbations, allowing differentiation between nonspecific cytotoxic responses and genuine mechanistic activity.

Predictive performance across assays was strongly influenced by the density of informative training signal, particularly the number of active compounds available for model learning (Figure 1). Endpoints with larger active fractions, such as cytochrome P450 antagonism assays, achieved the highest precision–recall scores and sensitivities because the models could learn more distinct structure–activity patterns. In contrast, assays with sparse active compounds, such as nuclear receptor or transcription factor pathways, exhibited weaker or less stable models, reflecting limited exposure to positive examples during training. This dependence on active compound size is consistent with observations from prior Tox21 and related HTS challenges, where endpoints with very low active fractions often yield weak or unstable models, while more balanced endpoints yield reproducible predictions^27^. Expanding the screening to larger compound libraries can partially mitigate this problem by increasing the absolute number of actives even in low-activity assays. However, scaling alone rarely resolves the challenge for inherently sparse biological targets. In such cases, improvements may rely more on leveraging complementary available information rather than simply increasing sample size. Multi-task prediction and transfer learning frameworks could offer one such solution, enabling models to learn shared patterns across related assays. By jointly modeling related endpoints, these approaches can capture overlapping molecular mechanisms and enhance stability and generalization without requiring exhaustive new experiments, particularly relevant for receptor and transcription factor assays where activity is context-specific and limited.

Beyond performance metrics, interpretability remains essential for the acceptance of machine learning models in regulatory toxicology. Black-box models can achieve high predictive accuracy but often fail to provide mechanistic transparency, limiting their trustworthiness in decision-making contexts^28^. Grounding predictive models in biologically rich data provides a path to overcome this limitation. While this study focused on HTTr, the same principles apply to other New Approach Methodologies (NAMs), including proteomics, metabolomics, and advanced imaging, which capture complementary facets of chemical activity^29^. In this study, SHAP-based feature attributions revealed that both structural motifs and transcriptional programs contributed to predictions in ways consistent with known principles of ligand– receptor interaction, stress adaptation, and cell survival regulation. This mechanistic transparency bridges predictive modeling and biological inference, supporting both regulatory acceptance and hypothesis generation.

Despite promising results, several factors limit the broader applicability of the current framework. Transcriptomic responses are inherently context-dependent, varying with cell type, receptor expression, and metabolic competence. Consequently models trained on compounds in one cellular context may not transfer optimally to another. For example, hepatocyte-derived systems exhibit rich xenobiotic metabolism but may underrepresent receptor-mediated signaling pathways in endocrine tissues^30^. Conversely, reporter-based or immortalized lines can capture receptor signaling yet lack metabolic activation capacity. Expanding the modeling framework to include a broader panel of metabolically competent and physiologically complementary systems should therefore improve both predictive robustness and translational realism. Furthermore, most available datasets emphasize single-chemical, short-duration exposures under uniform culture conditions. In contrast, real-world toxicological risk reflects complex scenarios involving chemical mixtures, repeated or chronic dosing, and life stage–specific susceptibilities. Incorporating such factors into future experimental designs and modeling pipelines will better approximate human biology and enhance the translational value of computational predictions. Another consideration concerns the compression of transcriptomic dose–response information into engineered summary features. While this design choice stabilizes modeling and facilitates throughput across thousands of chemical–assay combinations, it inevitably simplifies dynamic and potency-related aspects of transcriptional responses. This means that fine-grained dose–response patterns are partially flattened into more general descriptors, and highlights the trade-off between scalability and resolution in data-driven toxicology, as the ability to scale while retaining mechanistic depth will ultimately determine how effectively such frameworks can inform future safety assessments. Future work could deepen this framework by exploring representation-learning approaches that directly model continuous dose– response trajectories, such as spline-based architectures or sequence models that capture temporal progression across concentrations. These methods offer a path to balance interpretability with higher-resolution mechanistic detail, potentially recovering subtleties that engineered summary features may overlook. In parallel, multi-omic extensions that integrate proteomic, metabolomic, or imaging-based modalities could further strengthen predictive capacity and provide a more holistic view of chemical action, advancing both mechanistic inference and regulatory applicability.

In summary, our results demonstrate that integrating chemical structure descriptors with transcriptomic-wide dose-response profiles across multiple cell-lines yields models that are both predictive and mechanistically interpretable across Tox21 assays. Predictive accuracy increased with the number of active compounds available for model training, reflecting the influence of class balance on learning stability. Nevertheless, integrating chemical structure with transcriptomic dose–response profiles provided complementary information that improved sensitivity and interpretability even under imbalance. This combined representation allows the models to leverage mechanistic features, such as coordinated gene expression programs and structural motifs linked to target engagement, rather than depending solely on class frequency, thereby supporting more biologically grounded predictions across diverse assays. This framework naturally extends to additional cell lines and more elaborate experimental approaches, including multi-omics, longer exposure durations, repeated dosing, and mixtures. In doing so, this study contributes to the evolution of computational toxicology toward new approach methodologies that are transparent, human-relevant, and mechanistically grounded.

## Methods

### ASSAY RETRIEVAL AND SELECTION

HTS data were extracted from the U.S. EPA’s ToxCast in vitro screening database (InvitroDB v4.2) using a local MySQL server and custom SQL query. For each Tox21 assay component endpoint, a binary activity call was generated for all tested chemicals in the corresponding assays. Briefly, records containing continuous hit-call values (hitc) and fit category values (fitc) were filtered to retain only those with reliable values corresponding to strong actives or inactives, as per the InvitroDB v4.2 documentation. These outcomes were subsequently linked with sample- and chemical-level annotations to associate each endpoint outcome with their corresponding distributed structure-searchable toxicity substance identifier (DTXSID). The resulting datasets were filtered for relevance, first by excluding files containing channel-specific readouts (ch1/ch2), follow-up studies, real-time measurements, and autofluorescence data that do not represent primary biological activity. Compounds flagged as cytotoxic in corresponding Tox21 viability assays were subsequently removed to minimize the influence of nonspecific or confounded responses, and datasets with insufficient numbers of positive or negative samples (≤200 in either class) were excluded to ensure adequate data for downstream analyses (Supplementary Table 1). All code used throughout this manuscript is available on GitHub (https://github.com/gcattebeke/Paper_Mechanistically_Interpretable_Toxicity_Prediction), and an overview of the workflow can be found in Figure 4.

**Figure 4:**
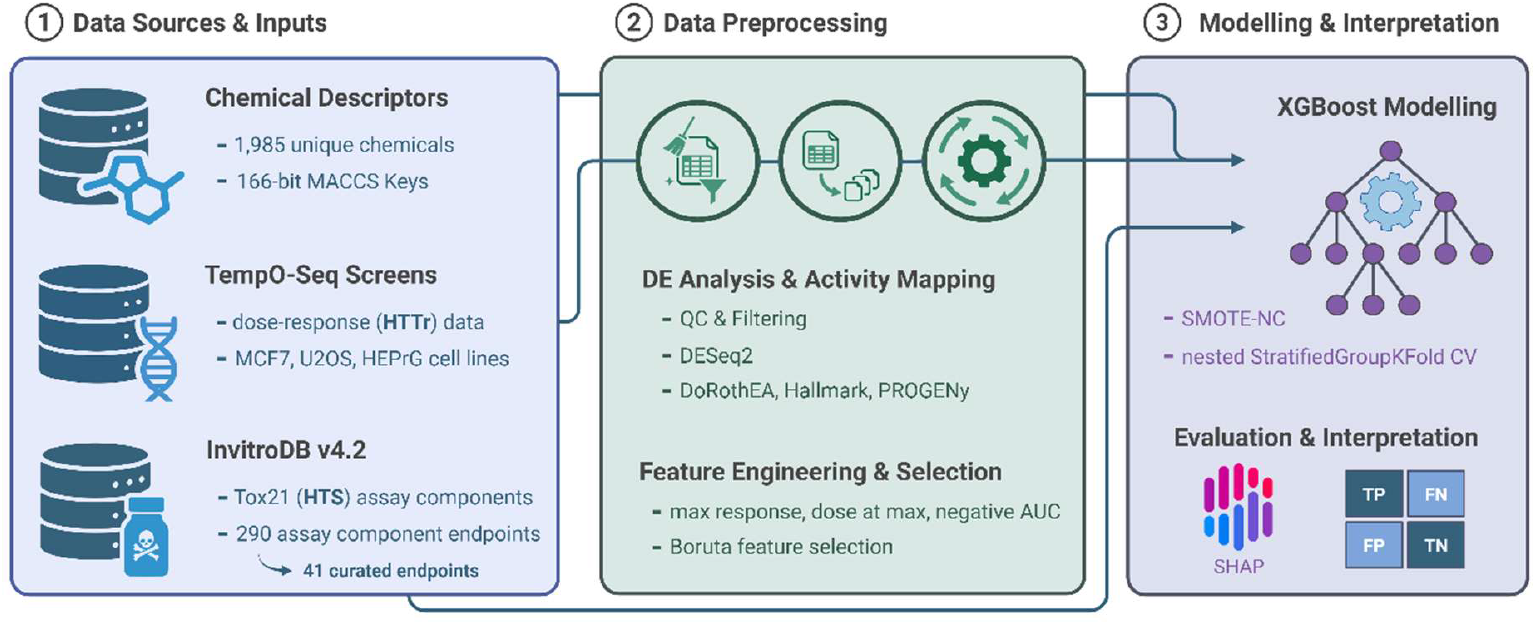
Overview of the modeling framework integrating chemical structure and transcriptomic dose–response data for mechanistically interpretable toxicity prediction. The workflow comprises three stages: (1) **Data sources and inputs** combine molecular fingerprints (166-bit MACCS keys) for 1,985 unique chemicals with high-throughput transcriptomic (HTTr) dose–response profiles from TempO-Seq screens (MCF-7, U-2 OS, and HepaRG cell lines) and curated assay outcomes from InvitroDB v4.2 (41 Tox21 endpoints). (2) **Data preprocessing** includes differential expression analysis using DESeq2 and activity mapping (DoRothEA, Hallmark, and PROGENy signatures) followed by feature engineering of dose–response summaries (maximum response, dose at maximum, and negative AUC) and Boruta-based feature selection to retain stable, informative transcriptomic descriptors. (3) **Modeling and interpretation** employ XGBoost classifiers trained under nested, compound-aware (StratifiedGroupKFold) cross-validation with SMOTE-NC balancing. Model outputs are interpreted through SHAP analysis to identify structural motifs and transcriptional programs driving assay activity.

### ENCODING CHEMICAL STRUCTURE

Chemical structure information was retrieved and processed to generate fingerprints for all compounds in this study. DTXSID identifiers were mapped to chemical annotations using the EPA’s CompTox Chemistry Dashboard batch search functionality. For compounds not initially identified using the standard chemical mapping files, manual searches were conducted to ensure a complete set. Using the SMILES (Simplified Molecular Input Line Entry System) representation, Molecular ACCess System (MACCS) keys were computed using the RDKit cheminformatics toolkit (v2024.03.5), resulting in 166-bit binary vectors encoding the presence or absence of specific predefined structural fragments within each molecule^31^. MACCS keys were selected because these predefined fragments enable direct SHAP attribution to recognizable chemical motifs, whereas circular or hashed fingerprints generate high-dimensional representations that hinder mechanistic interpretation. Compounds with invalid SMILES representations were flagged and excluded.

### DIFFERENTIAL GENE EXPRESSION AND ACTIVITY MAPPING

Probe-level count matrices from TempO-Seq high-throughput transcriptomic screens were retrieved from the NCBI Gene Expression Omnibus and the EPA’s EDAP Clowder repository for three human cell lines: MCF-7 breast cancer epithelial cells (GSE272548; 1,751 chemicals, 6 h exposure, 8 concentrations), U-2 OS osteosarcoma epithelial cells (EPA EDAP Clowder repository, dataset COMPTOX_Public; 1,201 chemicals, 24 h exposure, 8 concentrations), and differentiated HepaRG hepatocyte-like cells (GSE284321; 1,201 chemicals, 24 h exposure, 8 concentrations). All data were processed in R (v4.3.3). Probe-level counts were first filtered to exclude probes with low overall expression (mean count < 5 across all samples within each respective batch group). Samples failing quality metrics as per provided metadata annotations were omitted before starting the analysis using DESeq2 (v1.42.1). Specifically, samples were excluded if they failed one or more quality control criteria, including low numbers of mapped reads, low fraction of uniquely mapped reads, insufficient probe coverage, abnormal signal distribution, or high Gini coefficient. Additionally, assay wells flagged for cytotoxicity in the paired viability screens (based on normalized cell count and apoptosis/membrane-integrity markers) were removed from downstream analyses. For each chemical, every nonzero concentration was contrasted against its vehicle control to estimate log2 fold changes (L2FC), accounting for batch effects in the DESeq2 design formula. L2FC estimates were stabilized using adaptive shrinkage (ashr) and adjusted for multiple testing via the Benjamini–Hochberg procedure^32,33^. Genes represented by multiple probes were collapsed at the gene level by selecting for each gene and concentration, the probe with the largest absolute L2FC.

To translate gene-level responses into interpretable biological activities, we generated per-sample enrichment signatures across multiple biological layers. Single-sample gene set enrichment analysis (ssGSEA) was performed using the MSigDB Hallmark (H) and Curated (C2) collections, excluding gene sets with fewer than five genes and applying internal normalization (ssgsea.norm = TRUE). Transcription factor activities were inferred using DoRothEA regulons (confidence A–C) with decoupleR (v2.8.0), integrating univariate, multivariate, and weighted-sum statistics into a consensus activity score. Lastly, pathway-level activities were estimated with PROGENy (scale = TRUE, z_scores = TRUE) using the top 5,000 most responsive genes.

### DOSE RESPONSE AGGREGATION AND DATA INTEGRATION

To integrate multi-dose transcriptional activity profiles with single-endpoint assay outcomes, we summarized each chemical’s dose–response profile into compact, biologically meaningful features. For each unique combination of DTXSID, EPA sample ID, and cell type, dose–response curves were processed to derive three summary metrics per HTTr signature: (i) the maximum response, defined as the median of the two highest absolute values; (ii) the normalized dose level at which the absolute maximum response occurred; and (iii) the area under the curve (AUC) for negative responses, calculated using the trapezoidal rule. These features act as proxies for the underlying transcriptional dose–response profile, capturing response intensity, chemical potency, and integrated transcriptional suppression, and were selected for computational efficiency and scalability. For interpretability, AUC values were sign-flipped such that larger values represent stronger transcriptional suppression. These aggregated features were linked to Tox21 assay outcomes via shared DTXSID identifiers, and MACCS structural fingerprints were subsequently appended. The resulting feature matrix combines chemical structure information with quantitative transcriptional dose–response descriptors for predicting *in vitro* toxicity endpoints.

### FEATURE SELECTION

The integrated feature matrix comprised 16,821 transcriptomic dose–response summary features and 166-bit MACCS structural fingerprints, covering 1,985 unique chemicals across 41 assay endpoints. Because MACCS keys already provide a compact, predefined representation of molecular fragments, feature selection was applied only to the dose–response summaries. Feature importance was assessed using the Boruta algorithm (BorutaPy v0.4.3) with a Random Forest classifier (scikit-learn v1.5.2). The Boruta algorithm is a wrapper method that iteratively compares each real feature’s importance against permuted “shadow” copies, retaining only those that consistently exceed the maximum importance observed among all shadow features across iterations^15^.

To ensure robustness and mitigate class imbalance between active and inactive compounds (positives and negatives, respectively), negative samples were randomly undersampled to match the number of positives where possible. This procedure was repeated five times with different random seeds, so Boruta could evaluate features under multiple balanced views of the training data rather than a single draw. Stable features were defined by consensus, retaining only those identified in at least 60% of the runs. A 60% (3/5 runs) consensus threshold was selected because it balances stability with sensitivity. The resulting matrix, composed of this stable subset of dose–response features together with the MACCS fingerprints, served as the final input for downstream model training, providing a balanced representation of chemical structure and biologically derived activity patterns.

### CLASSIFICATION MODELLING AND PERFORMANCE ASSESSEMENT

Supervised binary classifiers were trained to predict each Tox21 assay endpoint from the integrated chemical and transcriptional feature space, ensuring that evaluation folds remained independent at the chemical level. To prevent compound-level information leakage, we implemented a nested, stratified, group-aware cross-validation workflow in which both the outer performance estimation and the inner hyperparameter selection treated each compound (DTXSID) as a grouping factor. In the outer loop, data were split into three folds using StratifiedGroupKFold, with one fold held out for unbiased model evaluation and the remaining folds used for training and validation. To address class imbalance between active and inactive compounds, we applied the Synthetic Minority Over-sampling Technique for Nominal and Continuous features (SMOTE-NC, imbalanced-learn v0.13.0) within the training folds. MACCS fingerprints were treated as categorical (nominal) features, whereas transcriptomic dose–response summaries were treated as continuous.

Gradient-boosted decision trees (XGBoost v2.1.1) were used as the predictive modeling framework, as they are known to deliver strong performance on structured, high-dimensional datasets. Within each inner cross-validation loop, hyperparameters including tree depth, learning rate, number of estimators, column subsampling, and L1/L2 regularization were optimized using RandomizedSearchCV across 50 candidate parameter sets, with mean average precision as the selection criterion. DTXSID-based grouping was consistently enforced during tuning to avoid chemical overlap between folds and thus to prevent compound-level leakage across folds. Predictive performance was evaluated using both (i) threshold-independent metrics including Area Under the Receiver Operating Characteristic Curve (AUROC), Area Under the Precision–Recall Curve (AUPRC), average precision, Brier score, and log loss, and (ii) threshold-dependent metrics including accuracy, F1-score, precision, recall, and specificity (Table 1). For each outer fold, the optimal classification threshold was selected by maximizing the F1-score on the precision–recall curve, and a single global threshold was subsequently derived from aggregated out-of-fold predictions.

To interpret model behavior, SHapley Additive exPlanations (SHAP) analysis was applied to the best-performing XGBoost model in each outer fold. A tree-based SHAP explainer was fitted using the SMOTE-NC–resampled training data as background, enabling the computation of local (sample-level) and global (feature-level) contributions to model predictions. Global importance was summarized per fold as the mean absolute SHAP value for each feature, providing interpretable insight into the structural and transcriptional drivers of assay activity.

## Data Availability

All data supporting this study are publicly available or included with the article and its Supplementary Information. TempO-Seq count matrices and sample metadata were downloaded from the NCBI Gene Expression Omnibus (GEO) under accessions GSE272548 (MCF-7) and GSE284321 (HepaRG), and from the EPA EDAP Clowder repository (collection: COMPTOX_Public) for U-2 OS. Curated HTS assay outcomes were obtained from U.S. EPA ToxCast/InvitroDB v4.2. The filtered per-endpoint table as extracted from the InVitroDB and used in this study, including AEIDs and activity hit calls, are available on GitHub (https://github.com/gcattebeke/Paper_Mechanistically_Interpretable_Toxicity_Prediction).

## Code Availability

All code used to reproduce data extraction, preprocessing, feature engineering, model training, evaluation, result analysis, and figure generation is publicly available on Github (https://github.com/gcattebeke/Paper_Mechanistically_Interpretable_Toxicity_Prediction) and Zenodo.

## Ethics Declarations

No human or animal subjects were directly involved in this study. The analyses were performed exclusively on publicly available transcriptomic datasets from the NCBI Gene Expression Omnibus and the EPA’s EDAP Clowder repository, as well as publicly released ToxCast/InvitroDB v4.2 assay data.

