## Supplementary Information for "Mechanistically Interpretable Toxicity Prediction Through Multimodal Integration of Structure and Transcriptomics"

This document contains supplementary information for the paper *"Mechanistically Interpretable Toxicity Prediction Through Multimodal Integration of Structure and Transcriptomics"*.

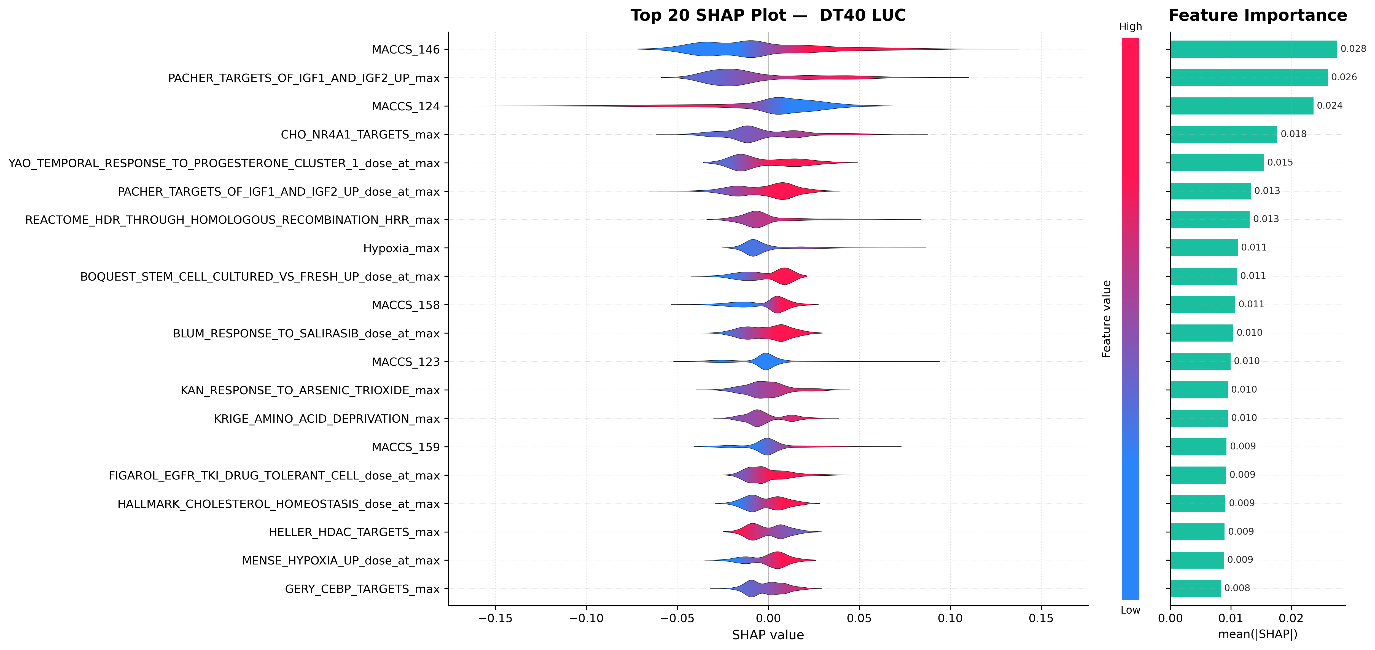

**Supplementary Figure 1: SHAP value analysis of the luciferase reporter assay in DT40 cells .** The violin plots (left) display the distribution of SHAP values for the top 20 most important features, ordered by mean absolute SHAP value (right). Each violin's width represents the density of SHAP values across all samples, with the x-axis indicating feature contribution magnitude and direction. The color gradient within violins encodes the normalized feature value revealing feature-prediction relationships.

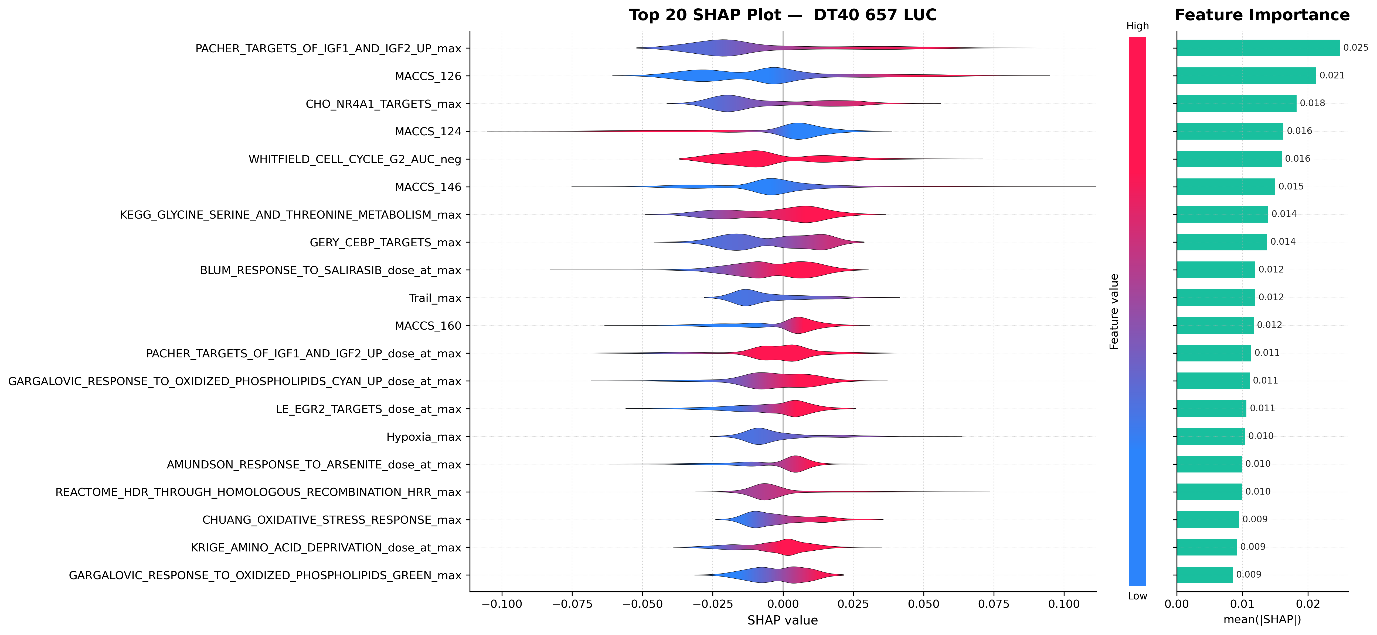

**Supplementary Figure 2: SHAP value analysis of the luciferase reporter assay in DT40 657 cells .** The violin plots (left) display the distribution of SHAP values for the top 20 most important features, ordered by mean absolute SHAP value (right). Each violin's width represents the density of SHAP values across all samples, with the x-axis indicating feature contribution magnitude and direction. The color gradient within violins encodes the normalized feature value revealing feature-prediction relationships.

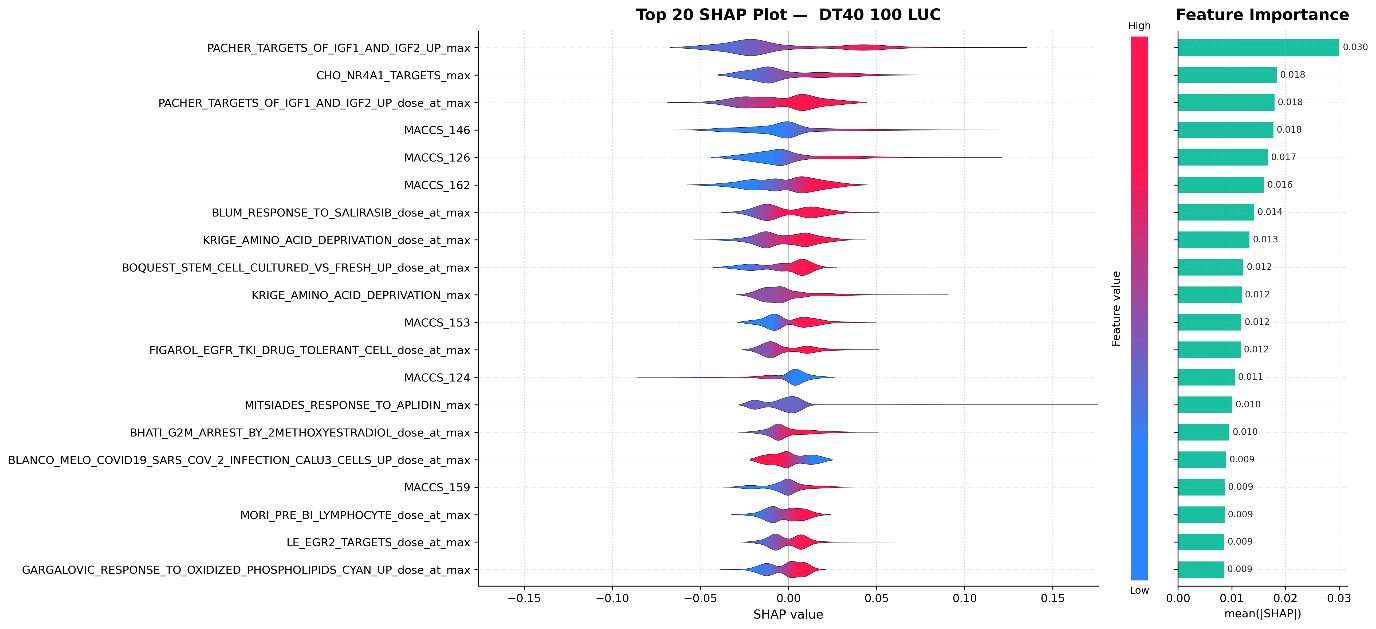

**Supplementary Figure 3: SHAP value analysis of the luciferase reporter assay in DT40 100 cells .** The violin plots (left) display the distribution of SHAP values for the top 20 most important features, ordered by mean absolute SHAP value (right). Each violin's width represents the density of SHAP values across all samples, with the x-axis indicating feature contribution magnitude and direction. The color gradient within violins encodes the normalized feature value revealing feature-prediction relationships.

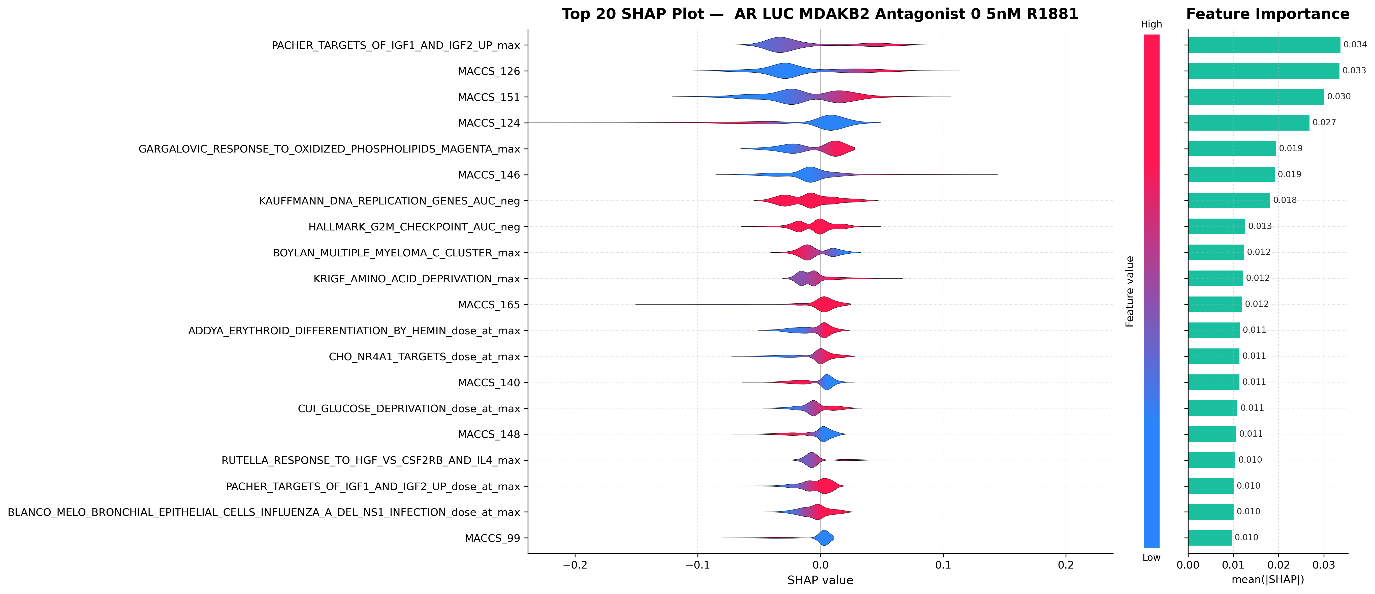

Supplementary Figure 4: **SHAP value analysis of the androgen receptor luciferase MDAKB2 antagonist assay in presence of 0.5nM R1881.** The violin plots (left) display the distribution of SHAP values for the top 20 most important features, ordered by mean absolute SHAP value (right). Each violin's width represents the density of SHAP values across all samples, with the x-axis indicating feature contribution magnitude and direction. The color gradient within violins encodes the normalized feature value revealing feature-prediction relationships.

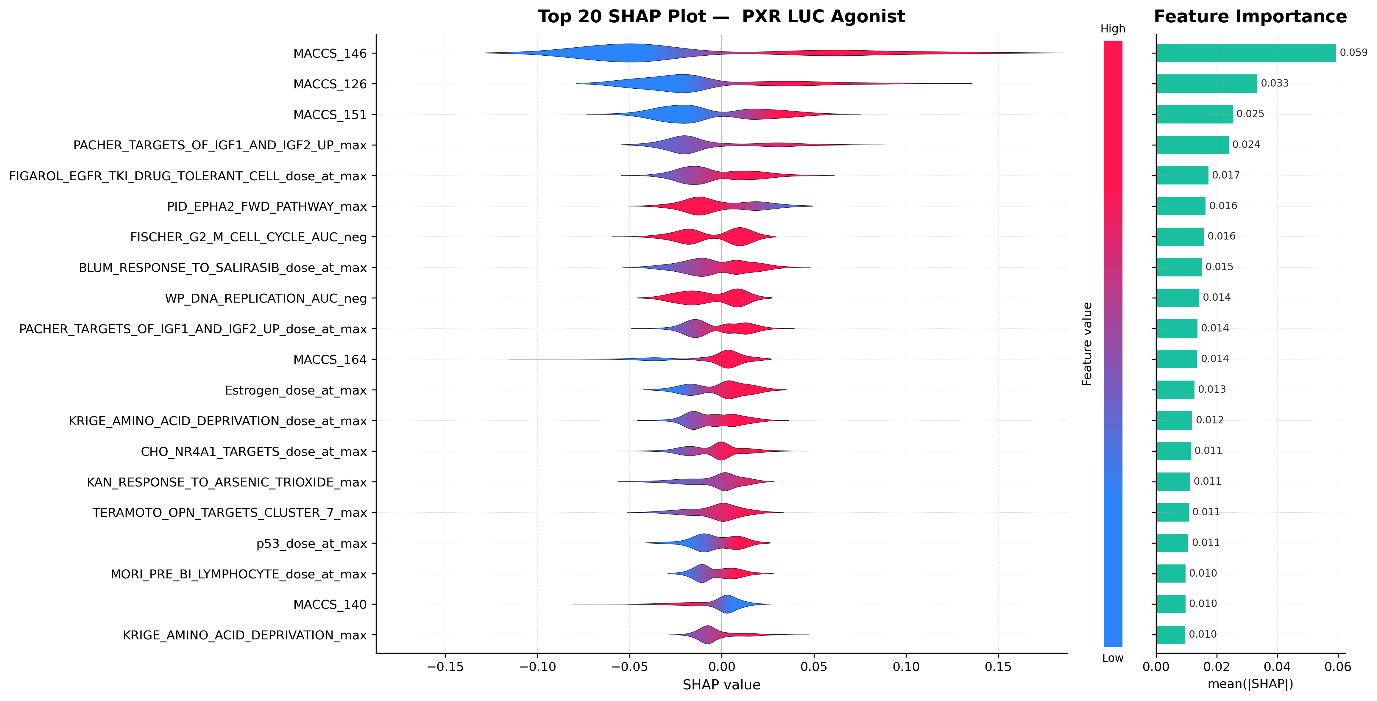

**Supplementary Figure 5:** **SHAP value analysis of the pregnane X receptor luciferase agonist assay.** The violin plots (left) display the distribution of SHAP values for the top 20 most important features, ordered by mean absolute SHAP value (right). Each violin's width represents the density of SHAP values across all samples, with the x-axis indicating feature contribution magnitude and direction. The color gradient within violins encodes the normalized feature value revealing feature-prediction relationships.

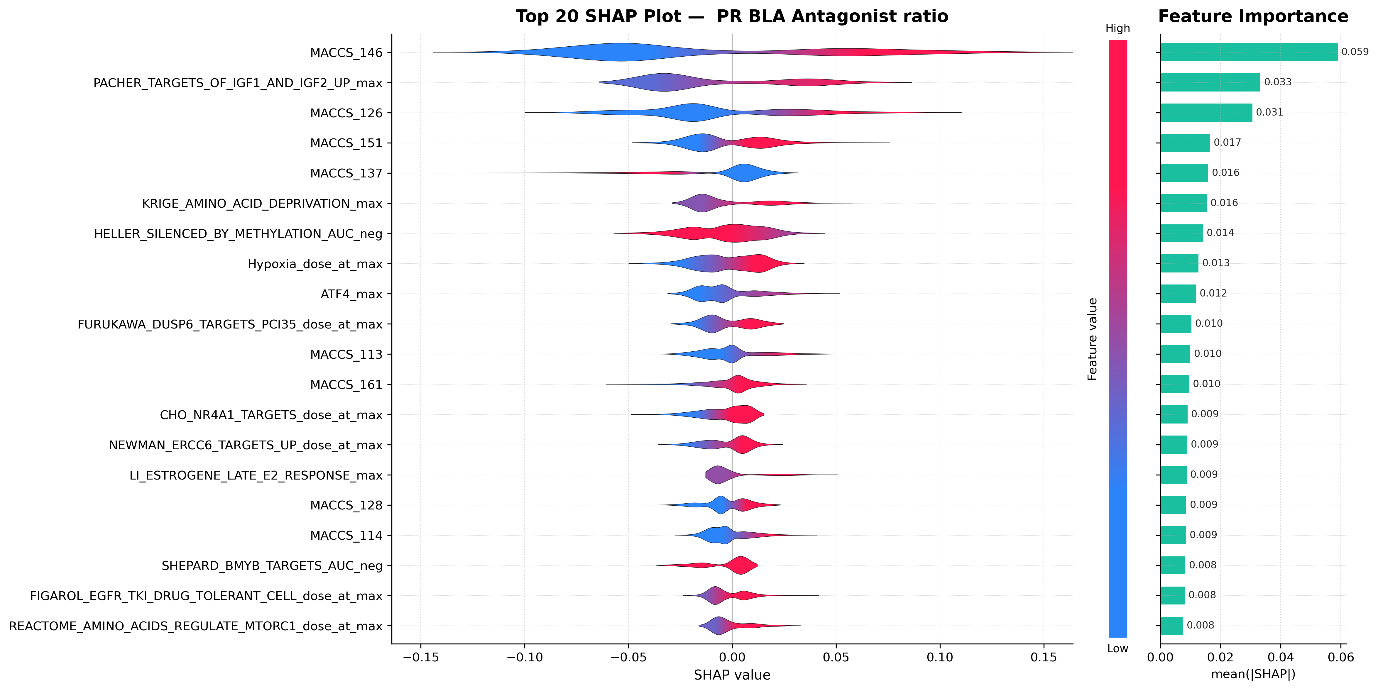

**Supplementary Figure 6: SHAP value analysis of the progesterone reporter beta lactamase assay.** The violin plots (left) display the distribution of SHAP values for the top 20 most important features, ordered by mean absolute SHAP value (right). Each violin's width represents the density of SHAP values across all samples, with the x-axis indicating feature contribution magnitude and direction. The color gradient within violins encodes the normalized feature value revealing feature-prediction relationships.

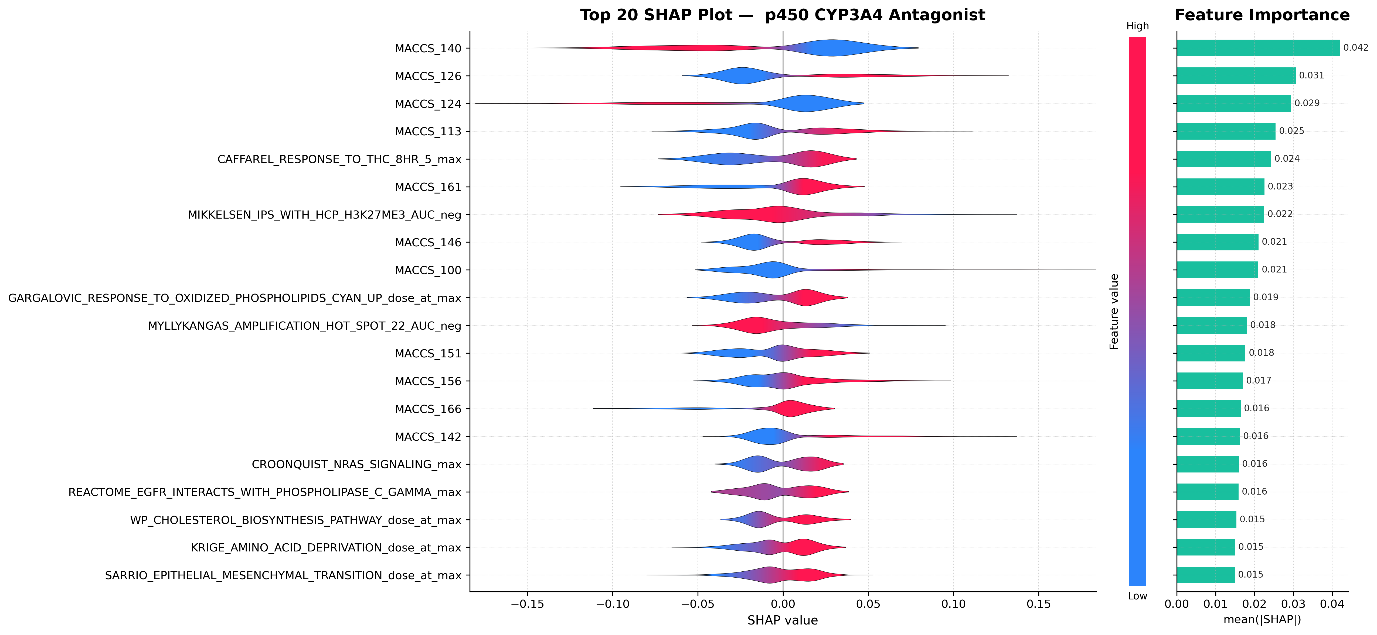

**Supplementary Figure 7: SHAP value analysis of the CYP3A4 Antagonist assay.** The violin plots (left) display the distribution of SHAP values for the top 20 most important features, ordered by mean absolute SHAP value (right). Each violin's width represents the density of SHAP values across all samples, with the x-axis indicating feature contribution magnitude and direction. The color gradient within violins encodes the normalized feature value revealing feature-prediction relationships

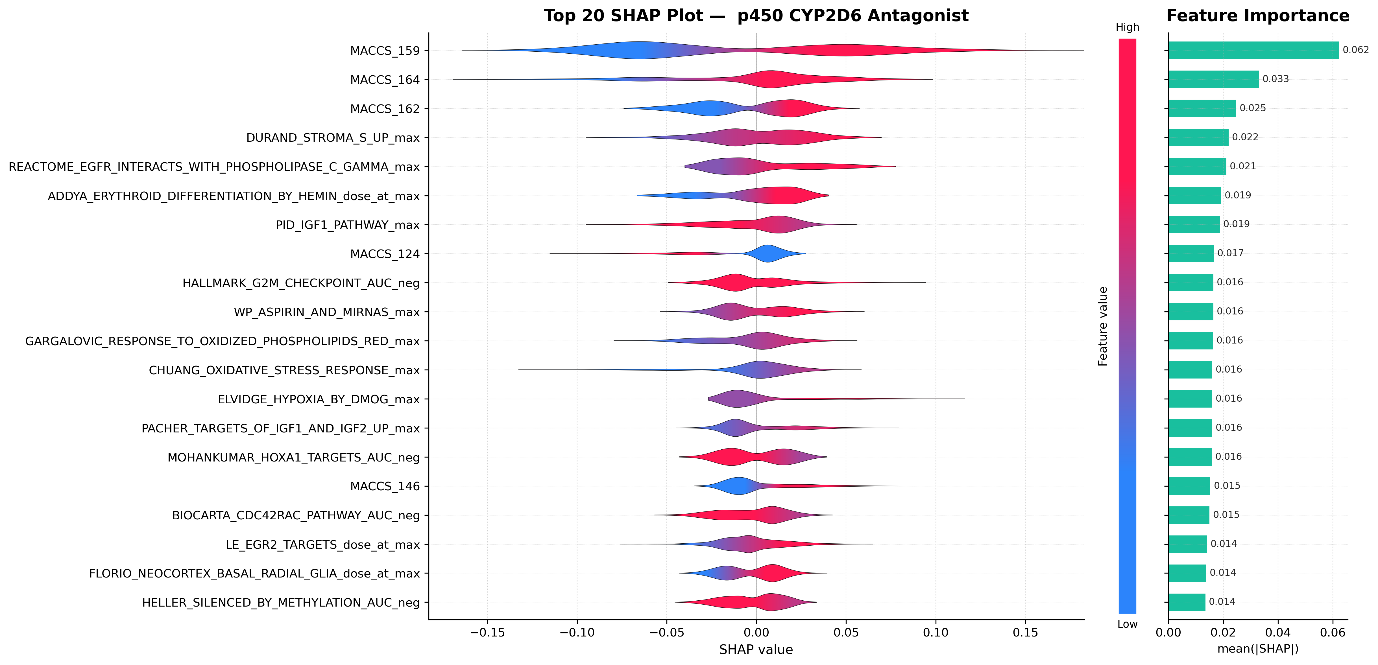

**Supplementary Figure 8: SHAP value analysis of the CYP2D6 Antagonist assay.** The violin plots (left) display the distribution of SHAP values for the top 20 most important features, ordered by mean absolute SHAP value (right). Each violin's width represents the density of SHAP values across all samples, with the x-axis indicating feature contribution magnitude and direction. The color gradient within violins encodes the normalized feature value revealing feature-prediction relationships.

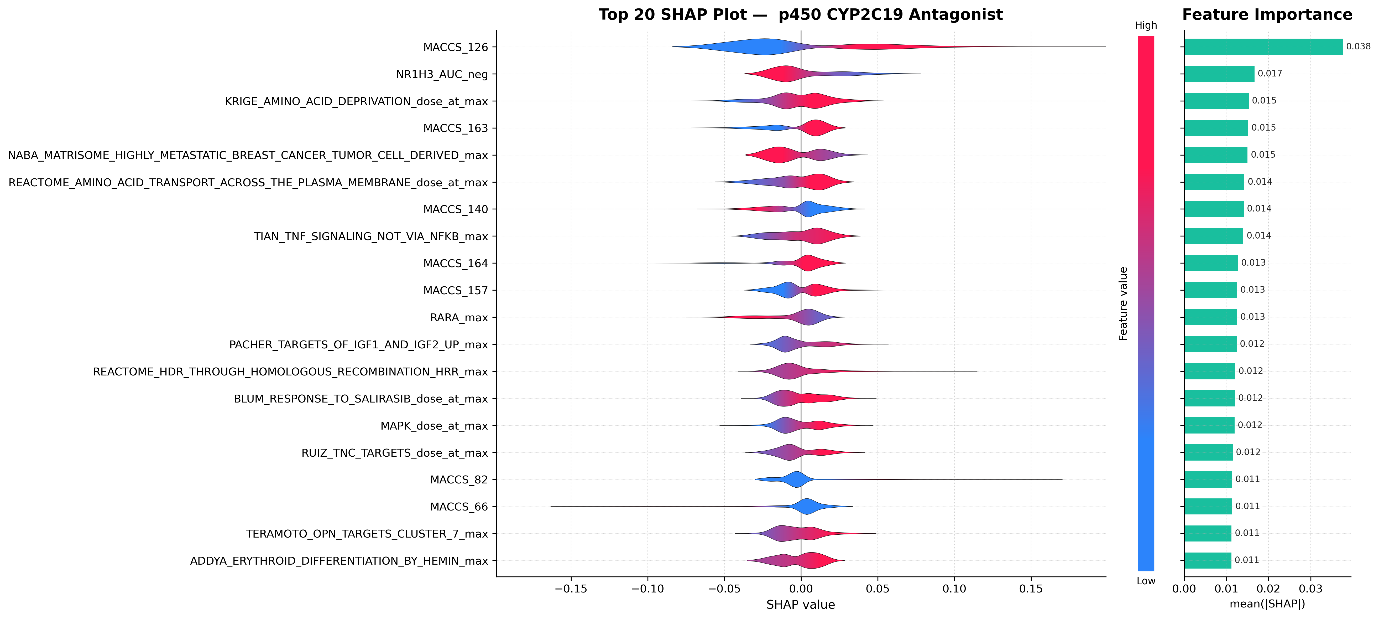

**Supplementary Figure 9: SHAP value analysis of the CYP2C19 Antagonist assay.** The violin plots (left) display the distribution of SHAP values for the top 20 most important features, ordered by mean absolute SHAP value (right). Each violin's width represents the density of SHAP values across all samples, with the x-axis indicating feature contribution magnitude and direction. The color gradient within violins encodes the normalized feature value revealing feature-prediction relationships.

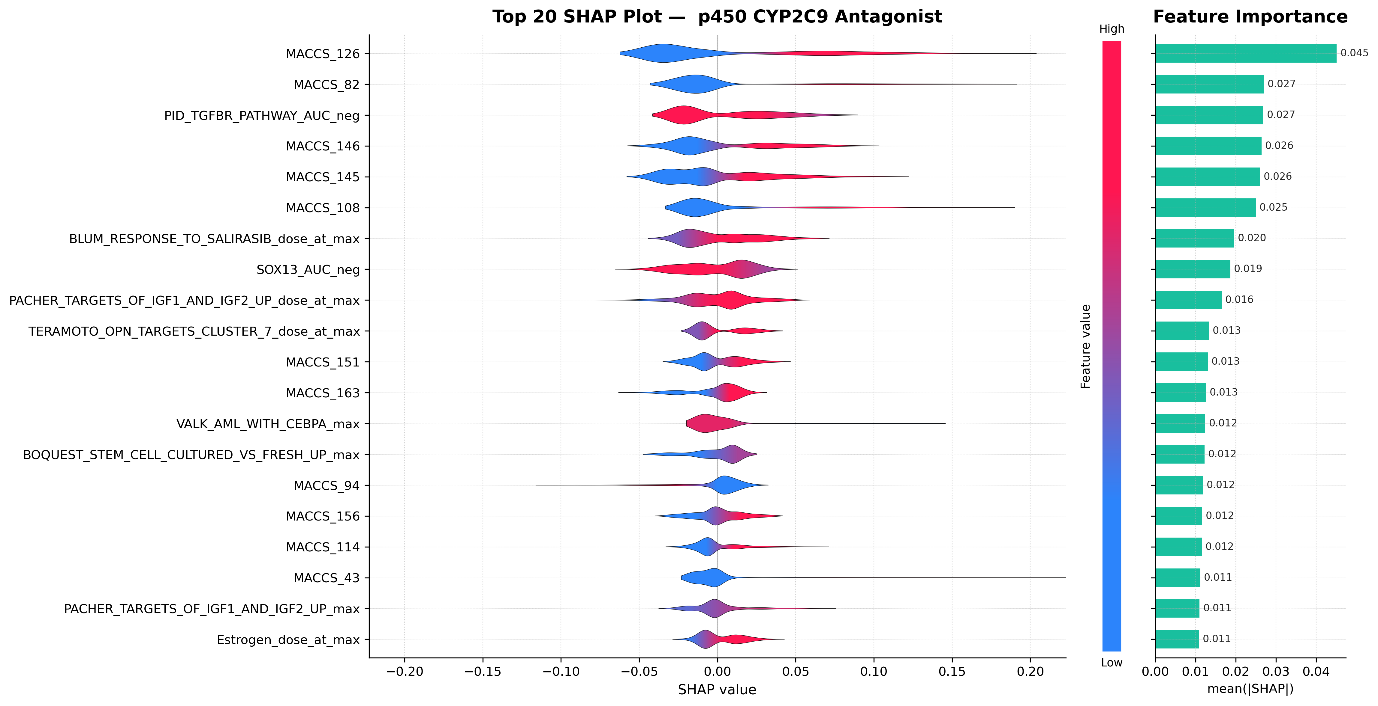

**Supplementary Figure 10: SHAP value analysis of the CYP2C9 Antagonist assay.** The violin plots (left) display the distribution of SHAP values for the top 20 most important features, ordered by mean absolute SHAP value (right). Each violin's width represents the density of SHAP values across all samples, with the x-axis indicating feature contribution magnitude and direction. The color gradient within violins encodes the normalized feature value revealing feature-prediction relationships.

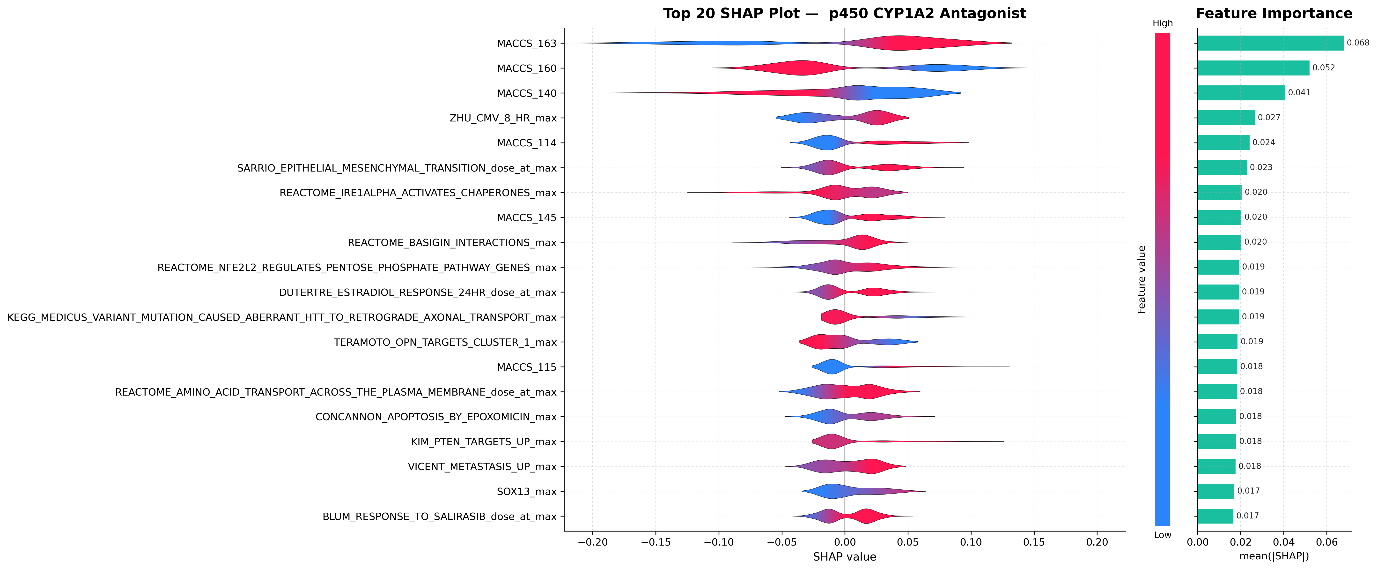

**Supplementary Figure 11: SHAP value analysis of the CYP1A2 Antagonist assay.** The violin plots (left) display the distribution of SHAP values for the top 20 most important features, ordered by mean absolute SHAP value (right). Each violin's width represents the density of SHAP values across all samples, with the x-axis indicating feature contribution magnitude and direction. The color gradient within violins encodes the normalized feature value revealing feature-prediction relationships.

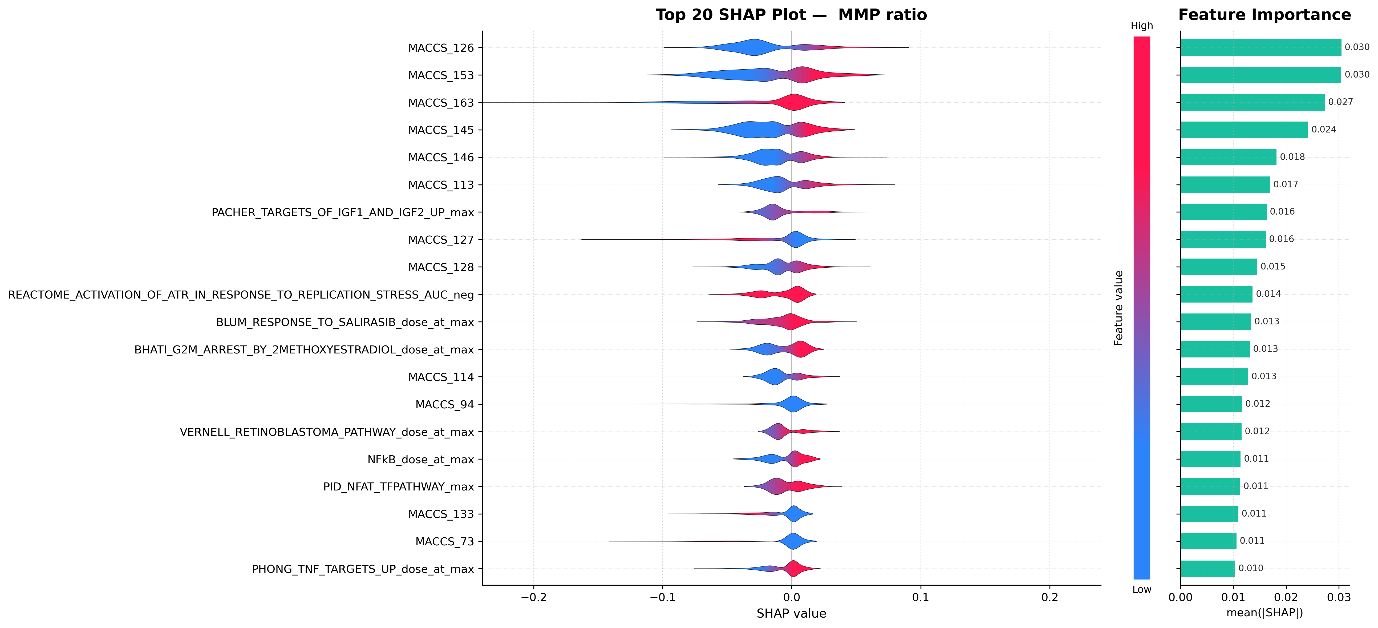

**Supplementary Figure 12:** **SHAP value analysis of the mitochondrial membrane potential assay.** The violin plots (left) display the distribution of SHAP values for the top 20 most important features, ordered by mean absolute SHAP value (right). Each violin's width represents the density of SHAP values across all samples, with the x-axis indicating feature contribution magnitude and direction. The color gradient within violins encodes the normalized feature value revealing feature-prediction relationships.

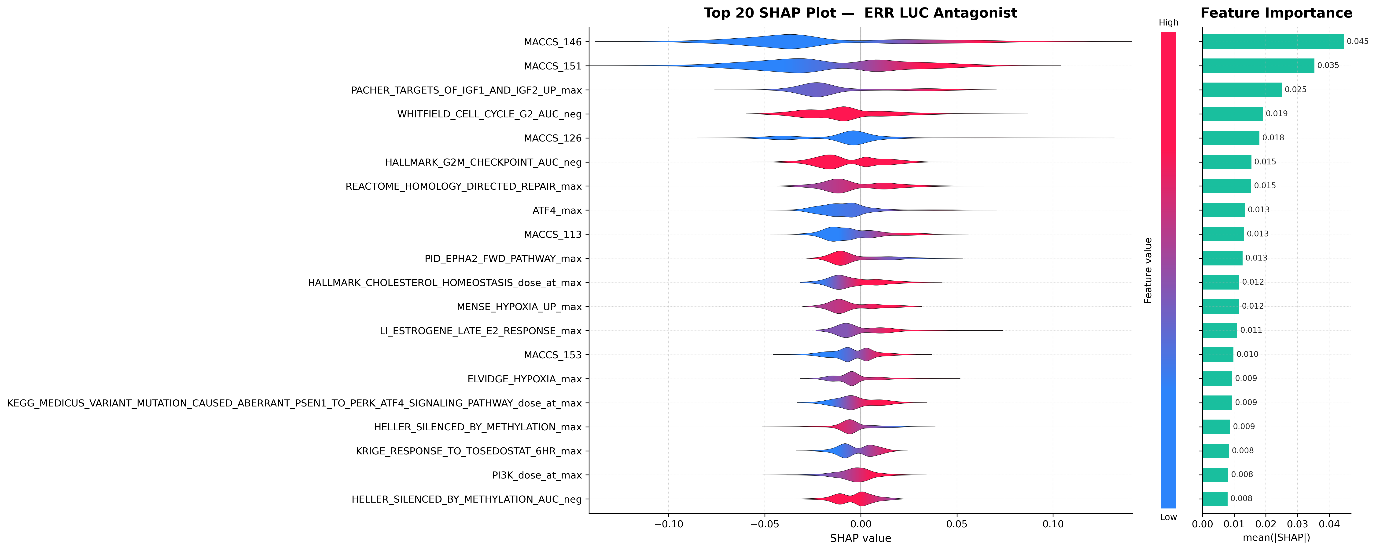

**Supplementary Figure 13:** **SHAP value analysis of the estrogen related receptor luciferase assay.** The violin plots (left) display the distribution of SHAP values for the top 20 most important features, ordered by mean absolute SHAP value (right). Each violin's width represents the density of SHAP values across all samples, with the x-axis indicating feature contribution magnitude and direction. The color gradient within violins encodes the normalized feature value revealing feature-prediction relationships.

**Supplementary Table 1: Comprehensive filtering characteristics of Tox21 assay endpoints showing exclusion criteria, sample counts, and final modeling dataset characteristics.** Assays underwent sequential filtering: (1) removal of technical replicates and unsuitable assay types; (2) cytotoxicity-based filtering using viability counterscreens; (3) minimum class size requirements (>200 per class). Final Total shows the number of chemical-cell type bags available for modeling after integration with HTTr transcriptomic features. HTTr Match Rate indicates the percentage of assay chemicals with available transcriptomic profiles. AEID: Assay Endpoint ID

| AEID | Assay Name | Exclusion Reason | Initial Total | Initial Positive | | Initial Negative | | Toxic Removed | | Final Total | Final Positive | Final Negative | | HTTr Match Rate (%) |
| --- | --- | --- | --- | --- | --- | --- | --- | --- | --- | --- | --- | --- | --- | --- |
| 2210 | TOX21_AChE_Colorimetric_Antagonist | Included | 6894 | 205 | | 6689 | |  | | 3625 | 153 | 3472 | | 26.3 |
| 806 | TOX21_AhR_LUC_Agonist | Included | 5949 | 244 | | 5705 | | 150 | | 2833 | 135 | 2698 | | 24.2 |
| 1846 | TOX21_AP1_BLA_Agonist_ratio | Included | 4971 | 394 | | 4577 | | 127 | | 2741 | 124 | 2617 | | 27.7 |
| 761 | TOX21_AR_BLA_Agonist_ratio | Included | 4870 | 214 | | 4656 | |  | | 2450 | 85 | 2365 | | 24.9 |
| 762 | TOX21_AR_BLA_Antagonist_ratio | Included | 5165 | 935 | | 4230 | | 428 | | 2352 | 371 | 1981 | | 24.6 |
| 1816 | TOX21_AR_LUC_MDAKB2_Antagonist_0.5nM_R1881 | Included | 4530 | | 1072 | | 3458 | | 255 | 2636 | 617 | 2019 | | 30.2 |
| 765 | TOX21_AR_LUC_MDAKB2_Antagonist_10nM_R1881 | Included | 5071 | 606 | | 4465 | | 331 | | 2552 | 225 | 2327 | | 26.7 |
| 1110 | TOX21_ARE_BLA_Agonist_ratio | Included | 4030 | 489 | | 3541 | | 51 | | 2269 | 293 | 1976 | | 27.9 |
| 767 | TOX21_Aromatase_LUC_Antagonist | Included | 4723 | 712 | | 4011 | | 493 | | 2080 | 222 | 1858 | | 23.8 |
| 2047 | TOX21_CAR_LUC_Agonist | Included | 6001 | 262 | | 5739 | | 107 | | 2928 | 228 | 2700 | | 25.1 |
| 2130 | TOX21_DT40_100_LUC | Included | 5344 | 2051 | | 3293 | |  | | 2911 | 1167 | 1744 | | 26.6 |
| 2131 | TOX21_DT40_657_LUC | Included | 5505 | 1980 | | 3525 | |  | | 2904 | 1146 | 1758 | | 26.1 |
| 1134 | TOX21_DT40_LUC | Included | 5192 | 2104 | | 3088 | |  | | 2753 | 1139 | 1614 | | 26.2 |
| 785 | TOX21_ERa_BLA_Agonist_ratio | Included | 7405 | 225 | | 7180 | |  | | 3390 | 128 | 3262 | | 22.8 |
| 786 | TOX21_ERa_BLA_Antagonist_ratio | Included | 4764 | 496 | | 4268 | | 130 | | 2249 | 269 | 1980 | | 23.8 |
| 788 | TOX21_ERa_LUC_VM7_Agonist | Included | 4576 | 446 | | 4130 | |  | | 2312 | 250 | 2062 | | 25 |
| 2053 | TOX21_ERa_LUC_VM7_Antagonist_0.1nM_E2 | Included | 4703 | | 561 | | 4142 | | 152 | 2612 | 299 | 2313 | | 28.3 |
| 789 | TOX21_ERa_LUC_VM7_Antagonist_0.5nM_E2 | Included | 4950 | | 451 | | 4499 | | 185 | 2446 | 224 | 2222 | | 25.1 |
| 2119 | TOX21_ERb_BLA_Antagonist_ratio | Included | 3952 | 867 | | 3085 | | 345 | | 2098 | 407 | 1691 | | 29.2 |
| 2057 | TOX21_ERR_LUC_Antagonist | Included | 4358 | 997 | | 3361 | | 329 | | 2334 | 525 | 1809 | | 28.5 |
| 1120 | TOX21_FXR_BLA_Antagonist_ratio | Included | 4634 | 558 | | 4076 | | 272 | | 2438 | 226 | 2212 | | 27.4 |
| 793 | TOX21_GR_BLA_Agonist_ratio | Included | 7311 | 283 | | 7028 | |  | | 3448 | 105 | 3343 | | 23.4 |
| 794 | TOX21_GR_BLA_Antagonist_ratio | Included | 5467 | 453 | | 5014 | | 110 | | 2694 | 223 | 2471 | | 24.8 |
| 2061 | TOX21_HDAC_LUC_Antagonist | Included | 6649 | 243 | | 6406 | | 41 | | 3463 | 124 | 3339 | | 26.3 |
| 3210 | TOX21_hERG_U2OS_Antagonist | Included | 5417 | 578 | | 4839 | |  | | 2929 | 272 | 2657 | | 27 |
| 1854 | TOX21_MMP_ratio | Included | 6916 | 790 | | 6126 | | 72 | | 3247 | 519 | 2728 | | 23.4 |
| 3188 | TOX21_p450_CYP1A2_Antagonist | Included | 4181 | 2254 | | 1927 | |  | | 2332 | 1436 | 896 | | 27.7 |
| 3186 | TOX21_p450_CYP2C19_Antagonist | Included | 5636 | 2094 | | 3542 | |  | | 3233 | 1505 | 1728 | | 28.5 |
| 3187 | TOX21_p450_CYP2C9_Antagonist | Included | 5748 | 2224 | | 3524 | |  | | 3050 | 1596 | 1454 | | 26.9 |
| 3185 | TOX21_p450_CYP2D6_Antagonist | Included | 4597 | 1809 | | 2788 | |  | | 2547 | 929 | 1618 | | 27.5 |
| 2544 | TOX21_p450_CYP3A4_Antagonist | Included | 4148 | 1422 | | 2726 | |  | | 2265 | 876 | 1389 | | 27.4 |
| 2070 | TOX21_PGC_ERR_LUC_Antagonist | Included | 4223 | 628 | | 3595 | | 255 | | 2329 | 324 | 2005 | | 29.2 |
| 1127 | TOX21_PPARg_BLA_Antagonist_ratio | Included | 4313 | 509 | | 3804 | | 192 | | 2460 | 260 | 2200 | | 29.1 |
| 2127 | TOX21_PR_BLA_Antagonist_ratio | Included | 3808 | 1224 | | 2584 | | 377 | | 2006 | 673 | 1333 | | 28.9 |
| 2363 | TOX21_PXR_LUC_Agonist | Included | 4147 | 1215 | | 2932 | | 252 | | 2177 | 701 | 1476 | | 27.7 |
| 1839 | TOX21_RAR_LUC_Antagonist | Included | 4124 | 493 | | 3631 | | 149 | | 2287 | 234 | 2053 | | 28 |
| 1661 | TOX21_RORg_LUC_CHO_Antagonist | Included | 4482 | 653 | | 3829 | | 305 | | 2461 | 263 | 2198 | | 29.3 |
| 2372 | TOX21_SBE_BLA_Antagonist_ratio | Included | 5267 | 599 | | 4668 | | 157 | | 2815 | 303 | 2512 | | 27.5 |
| 2108 | TOX21_SHH_3T3_GLI3_LUC_Antagonist | Included | 4153 | 1187 | | 2966 | | 555 | | 2131 | 444 | 1687 | | 29.3 |
| 804 | TOX21_TR_LUC_GH3_Antagonist | Included | 4075 | 1421 | | 2654 | | 973 | | 1631 | 256 | 1375 | | 26 |
| 2040 | TOX21_TSHR_HTRF_Agonist_ratio | Included | 5285 | 245 | | 5040 | |  | | 2914 | 201 | 2713 | | 27.4 |
| 769 | TOX21_AutoFluor_HEK293_Cell_blue | Autofluorescence | 4406 | 0 | | 4406 | |  | |  |  |  | |  |
| 770 | TOX21_AutoFluor_HEK293_Cell_green | Autofluorescence | 4877 | 0 | | 4877 | |  | |  |  |  | |  |
| 771 | TOX21_AutoFluor_HEK293_Cell_red | Autofluorescence | 4892 | 0 | | 4892 | |  | |  |  |  | |  |
| 772 | TOX21_AutoFluor_HEK293_Media_blue | Autofluorescence | 4407 | 0 | | 4407 | |  | |  |  |  | |  |
| 773 | TOX21_AutoFluor_HEK293_Media_green | Autofluorescence | 4749 | 0 | | 4749 | |  | |  |  |  | |  |
| 774 | TOX21_AutoFluor_HEK293_Media_red | Autofluorescence | 4790 | 0 | | 4790 | |  | |  |  |  | |  |
| 775 | TOX21_AutoFluor_HEPG2_Cell_blue | Autofluorescence | 4133 | 0 | | 4133 | |  | |  |  |  | |  |
| 776 | TOX21_AutoFluor_HEPG2_Cell_green | Autofluorescence | 3163 | 0 | | 3163 | |  | |  |  |  | |  |
| 777 | TOX21_AutoFluor_HEPG2_Cell_red | Autofluorescence | 3098 | 0 | | 3098 | |  | |  |  |  | |  |
| 778 | TOX21_AutoFluor_HEPG2_Media_blue | Autofluorescence | 4511 | 0 | | 4511 | |  | |  |  |  | |  |
| 779 | TOX21_AutoFluor_HEPG2_Media_green | Autofluorescence | 3181 | 0 | | 3181 | |  | |  |  |  | |  |
| 780 | TOX21_AutoFluor_HEPG2_Media_red | Autofluorescence | 4623 | 0 | | 4623 | |  | |  |  |  | |  |
| 1844 | TOX21_AP1_BLA_Agonist_ch1 | Channel-specific | 5673 | 338 | | 5335 | |  | |  |  |  | |  |
| 1845 | TOX21_AP1_BLA_Agonist_ch2 | Channel-specific | 4778 | 198 | | 4580 | |  | |  |  |  | |  |
| 759 | TOX21_AR_BLA_Agonist_ch1 | Channel-specific | 5939 | 52 | | 5887 | |  | |  |  |  | |  |
| 760 | TOX21_AR_BLA_Agonist_ch2 | Channel-specific | 5747 | 184 | | 5563 | |  | |  |  |  | |  |
| 1202 | TOX21_AR_BLA_Antagonist_ch1 | Channel-specific | 6231 | 40 | | 6191 | |  | |  |  |  | |  |
| 1203 | TOX21_AR_BLA_Antagonist_ch2 | Channel-specific | 5704 | 672 | | 5032 | |  | |  |  |  | |  |
| 1108 | TOX21_ARE_BLA_Agonist_ch1 | Channel-specific | 5032 | 198 | | 4834 | |  | |  |  |  | |  |
| 1109 | TOX21_ARE_BLA_Agonist_ch2 | Channel-specific | 4274 | 565 | | 3709 | |  | |  |  |  | |  |
| 783 | TOX21_ERa_BLA_Agonist_ch1 | Channel-specific | 5868 | 71 | | 5797 | |  | |  |  |  | |  |
| 784 | TOX21_ERa_BLA_Agonist_ch2 | Channel-specific | 7249 | 220 | | 7029 | |  | |  |  |  | |  |
| 1189 | TOX21_ERa_BLA_Antagonist_ch1 | Channel-specific | 5802 | 188 | | 5614 | |  | |  |  |  | |  |
| 1190 | TOX21_ERa_BLA_Antagonist_ch2 | Channel-specific | 5618 | 281 | | 5337 | |  | |  |  |  | |  |
| 2113 | TOX21_ERb_BLA_Agonist_ch1 | Channel-specific | 5158 | 40 | | 5118 | |  | |  |  |  | |  |
| 2114 | TOX21_ERb_BLA_Agonist_ch2 | Channel-specific | 5823 | 72 | | 5751 | |  | |  |  |  | |  |
| 2117 | TOX21_ERb_BLA_Antagonist_ch1 | Channel-specific | 5869 | 97 | | 5772 | |  | |  |  |  | |  |
| 2118 | TOX21_ERb_BLA_Antagonist_ch2 | Channel-specific | 4430 | 614 | | 3816 | |  | |  |  |  | |  |
| 1340 | TOX21_ESRE_BLA_Agonist_ch1 | Channel-specific | 4918 | 74 | | 4844 | |  | |  |  |  | |  |
| 1341 | TOX21_ESRE_BLA_Agonist_ch2 | Channel-specific | 6336 | 32 | | 6304 | |  | |  |  |  | |  |
| 1117 | TOX21_FXR_BLA_Agonist_ch1 | Channel-specific | 5252 | 31 | | 5221 | |  | |  |  |  | |  |
| 1118 | TOX21_FXR_BLA_Agonist_ch2 | Channel-specific | 6649 | 27 | | 6622 | |  | |  |  |  | |  |
| 1191 | TOX21_FXR_BLA_Antagonist_ch1 | Channel-specific | 4863 | 140 | | 4723 | |  | |  |  |  | |  |
| 1192 | TOX21_FXR_BLA_Antagonist_ch2 | Channel-specific | 4838 | 415 | | 4423 | |  | |  |  |  | |  |
| 791 | TOX21_GR_BLA_Agonist_ch1 | Channel-specific | 5413 | 241 | | 5172 | |  | |  |  |  | |  |
| 792 | TOX21_GR_BLA_Agonist_ch2 | Channel-specific | 6160 | 135 | | 6025 | |  | |  |  |  | |  |
| 1193 | TOX21_GR_BLA_Antagonist_ch1 | Channel-specific | 5875 | 11 | | 5864 | |  | |  |  |  | |  |
| 1194 | TOX21_GR_BLA_Antagonist_ch2 | Channel-specific | 5254 | 599 | | 4655 | |  | |  |  |  | |  |
| 1841 | TOX21_H2AX_HTRF_CHO_Agonist_ch1 | Channel-specific | 6168 | 0 | | 6168 | |  | |  |  |  | |  |
| 1842 | TOX21_H2AX_HTRF_CHO_Agonist_ch2 | Channel-specific | 6495 | 23 | | 6472 | |  | |  |  |  | |  |
| 2063 | TOX21_HRE_BLA_Agonist_ch1 | Channel-specific | 6459 | 308 | | 6151 | |  | |  |  |  | |  |
| 2064 | TOX21_HRE_BLA_Agonist_ch2 | Channel-specific | 7500 | 17 | | 7483 | |  | |  |  |  | |  |
| 1111 | TOX21_HSE_BLA_Agonist_ch1 | Channel-specific | 5654 | 173 | | 5481 | |  | |  |  |  | |  |
| 1112 | TOX21_HSE_BLA_Agonist_ch2 | Channel-specific | 6788 | 44 | | 6744 | |  | |  |  |  | |  |
| 1344 | TOX21_NFkB_BLA_agonist_ch1 | Channel-specific | 4986 | 195 | | 4791 | |  | |  |  |  | |  |
| 1345 | TOX21_NFkB_BLA_agonist_ch2 | Channel-specific | 7225 | 15 | | 7210 | |  | |  |  |  | |  |
| 1114 | TOX21_p53_BLA_p1_ch1 | Channel-specific | 5863 | 277 | | 5586 | |  | |  |  |  | |  |
| 1115 | TOX21_p53_BLA_p1_ch2 | Channel-specific | 7109 | 116 | | 6993 | |  | |  |  |  | |  |
| 1315 | TOX21_p53_BLA_p2_ch1 | Channel-specific | 2666 | 14 | | 2652 | |  | |  |  |  | |  |
| 1316 | TOX21_p53_BLA_p2_ch2 | Channel-specific | 3728 | 10 | | 3718 | |  | |  |  |  | |  |
| 1319 | TOX21_p53_BLA_p3_ch1 | Channel-specific | 3224 | 6 | | 3218 | |  | |  |  |  | |  |
| 1320 | TOX21_p53_BLA_p3_ch2 | Channel-specific | 3603 | 8 | | 3595 | |  | |  |  |  | |  |
| 1323 | TOX21_p53_BLA_p4_ch1 | Channel-specific | 3795 | 16 | | 3779 | |  | |  |  |  | |  |
| 1324 | TOX21_p53_BLA_p4_ch2 | Channel-specific | 4067 | 4 | | 4063 | |  | |  |  |  | |  |
| 1327 | TOX21_p53_BLA_p5_ch1 | Channel-specific | 3669 | 14 | | 3655 | |  | |  |  |  | |  |
| 1328 | TOX21_p53_BLA_p5_ch2 | Channel-specific | 4082 | 6 | | 4076 | |  | |  |  |  | |  |
| 1122 | TOX21_PPARd_BLA_Agonist_ch1 | Channel-specific | 5856 | 11 | | 5845 | |  | |  |  |  | |  |
| 1123 | TOX21_PPARd_BLA_Agonist_ch2 | Channel-specific | 6311 | 37 | | 6274 | |  | |  |  |  | |  |
| 1196 | TOX21_PPARd_BLA_Antagonist_ch1 | Channel-specific | 5514 | 81 | | 5433 | |  | |  |  |  | |  |
| 1197 | TOX21_PPARd_BLA_Antagonist_ch2 | Channel-specific | 4999 | 266 | | 4733 | |  | |  |  |  | |  |
| 800 | TOX21_PPARg_BLA_Agonist_ch1 | Channel-specific | 6604 | 33 | | 6571 | |  | |  |  |  | |  |
| 801 | TOX21_PPARg_BLA_Agonist_ch2 | Channel-specific | 5938 | 122 | | 5816 | |  | |  |  |  | |  |
| 1198 | TOX21_PPARg_BLA_Antagonist_ch1 | Channel-specific | 6238 | 111 | | 6127 | |  | |  |  |  | |  |
| 1199 | TOX21_PPARg_BLA_Antagonist_ch2 | Channel-specific | 4551 | 239 | | 4312 | |  | |  |  |  | |  |
| 2121 | TOX21_PR_BLA_Agonist_ch1 | Channel-specific | 4579 | 69 | | 4510 | |  | |  |  |  | |  |
| 2122 | TOX21_PR_BLA_Agonist_ch2 | Channel-specific | 5615 | 146 | | 5469 | |  | |  |  |  | |  |
| 2125 | TOX21_PR_BLA_Antagonist_ch1 | Channel-specific | 5343 | 84 | | 5259 | |  | |  |  |  | |  |
| 2126 | TOX21_PR_BLA_Antagonist_ch2 | Channel-specific | 4537 | 890 | | 3647 | |  | |  |  |  | |  |
| 2214 | TOX21_PR_BLA_Followup_Agonist_ch1 | Channel-specific | 134 | 0 | | 134 | |  | |  |  |  | |  |
| 2216 | TOX21_PR_BLA_Followup_Agonist_ch2 | Channel-specific | 142 | 21 | | 121 | |  | |  |  |  | |  |
| 2215 | TOX21_PR_BLA_Followup_Antagonist_ch1 | Channel-specific | 166 | 5 | | 161 | |  | |  |  |  | |  |
| 2217 | TOX21_PR_BLA_Followup_Antagonist_ch2 | Channel-specific | 158 | 4 | | 154 | |  | |  |  |  | |  |
| 1818 | TOX21_RXR_BLA_Agonist_ch1 | Channel-specific | 5121 | 4 | | 5117 | |  | |  |  |  | |  |
| 1819 | TOX21_RXR_BLA_Agonist_ch2 | Channel-specific | 4618 | 202 | | 4416 | |  | |  |  |  | |  |
| 2374 | TOX21_SBE_BLA_Agonist_ch1 | Channel-specific | 5435 | 1 | | 5434 | |  | |  |  |  | |  |
| 2375 | TOX21_SBE_BLA_Agonist_ch2 | Channel-specific | 7282 | 0 | | 7282 | |  | |  |  |  | |  |
| 2370 | TOX21_SBE_BLA_Antagonist_ch1 | Channel-specific | 5935 | 0 | | 5935 | |  | |  |  |  | |  |
| 2371 | TOX21_SBE_BLA_Antagonist_ch2 | Channel-specific | 5695 | 424 | | 5271 | |  | |  |  |  | |  |
| 2251 | TOX21_TR_RXR_BLA_Agonist_Followup_ch1 | Channel-specific | 46 | 0 | | 46 | |  | |  |  |  | |  |
| 2252 | TOX21_TR_RXR_BLA_Agonist_Followup_ch2 | Channel-specific | 58 | 4 | | 54 | |  | |  |  |  | |  |
| 2255 | TOX21_TR_RXR_BLA_Antagonist_Followup_ch1 | Channel-specific | 100 | 4 | | 96 | |  | |  |  |  | |  |
| 2256 | TOX21_TR_RXR_BLA_Antagonist_Followup_ch2 | Channel-specific | 69 | 0 | | 69 | |  | |  |  |  | |  |
| 2228 | TOX21_TRA_COA_Agonist_Followup_ch1 | Channel-specific | 72 | 0 | | 72 | |  | |  |  |  | |  |
| 2229 | TOX21_TRA_COA_Agonist_Followup_ch2 | Channel-specific | 66 | 8 | | 58 | |  | |  |  |  | |  |
| 2231 | TOX21_TRA_COA_Antagonist_Followup_ch1 | Channel-specific | 62 | 1 | | 61 | |  | |  |  |  | |  |
| 2232 | TOX21_TRA_COA_Antagonist_Followup_ch2 | Channel-specific | 116 | 4 | | 112 | |  | |  |  |  | |  |
| 2234 | TOX21_TRB_BLA_Agonist_Followup_ch1 | Channel-specific | 47 | 0 | | 47 | |  | |  |  |  | |  |
| 2235 | TOX21_TRB_BLA_Agonist_Followup_ch2 | Channel-specific | 22 | 0 | | 22 | |  | |  |  |  | |  |
| 2238 | TOX21_TRB_BLA_Antagonist_Followup_ch1 | Channel-specific | 90 | 3 | | 87 | |  | |  |  |  | |  |
| 2239 | TOX21_TRB_BLA_Antagonist_Followup_ch2 | Channel-specific | 73 | 1 | | 72 | |  | |  |  |  | |  |
| 2242 | TOX21_TRB_COA_Agonist_Followup_ch1 | Channel-specific | 71 | 0 | | 71 | |  | |  |  |  | |  |
| 2243 | TOX21_TRB_COA_Agonist_Followup_ch2 | Channel-specific | 66 | 9 | | 57 | |  | |  |  |  | |  |
| 2245 | TOX21_TRB_COA_Antagonist_Followup_ch1 | Channel-specific | 87 | 1 | | 86 | |  | |  |  |  | |  |
| 2246 | TOX21_TRB_COA_Antagonist_Followup_ch2 | Channel-specific | 114 | 7 | | 107 | |  | |  |  |  | |  |
| 2038 | TOX21_TSHR_HTRF_Agonist_ch1 | Channel-specific | 5990 | 263 | | 5727 | |  | |  |  |  | |  |
| 2039 | TOX21_TSHR_HTRF_Agonist_ch2 | Channel-specific | 6951 | 5 | | 6946 | |  | |  |  |  | |  |
| 2041 | TOX21_TSHR_HTRF_Antagonist_ch1 | Channel-specific | 6564 | 116 | | 6448 | |  | |  |  |  | |  |
| 2042 | TOX21_TSHR_HTRF_Antagonist_ch2 | Channel-specific | 6547 | 52 | | 6495 | |  | |  |  |  | |  |
| 2044 | TOX21_TSHR_wt_HTRF_Agonist_ch1 | Channel-specific | 6865 | 5 | | 6860 | |  | |  |  |  | |  |
| 2045 | TOX21_TSHR_wt_HTRF_Agonist_ch2 | Channel-specific | 7078 | 9 | | 7069 | |  | |  |  |  | |  |
| 1129 | TOX21_VDR_BLA_Agonist_ch1 | Channel-specific | 5437 | 1 | | 5436 | |  | |  |  |  | |  |
| 1130 | TOX21_VDR_BLA_Agonist_ch2 | Channel-specific | 5654 | 26 | | 5628 | |  | |  |  |  | |  |
| 1200 | TOX21_VDR_BLA_Antagonist_ch1 | Channel-specific | 5620 | 155 | | 5465 | |  | |  |  |  | |  |
| 1201 | TOX21_VDR_BLA_Antagonist_ch2 | Channel-specific | 4764 | 156 | | 4608 | |  | |  |  |  | |  |
| 2218 | TOX21_PR_BLA_Followup_Agonist_ratio | Follow-up assay | 194 | 33 | | 161 | |  | |  |  |  | |  |
| 2220 | TOX21_PR_BLA_Followup_Agonist_viability | Follow-up assay | 201 | 1 | | 200 | |  | |  |  |  | |  |
| 2219 | TOX21_PR_BLA_Followup_Antagonist_ratio | Follow-up assay | 125 | 110 | | 15 | |  | |  |  |  | |  |
| 2221 | TOX21_PR_BLA_Followup_Antagonist_viability | Follow-up assay | 202 | 0 | | 202 | |  | |  |  |  | |  |
| 2222 | TOX21_PR_LUC_Followup_Agonist | Follow-up assay | 171 | 45 | | 126 | |  | |  |  |  | |  |
| 2224 | TOX21_PR_LUC_Followup_Agonist_viability | Follow-up assay | 170 | 0 | | 170 | |  | |  |  |  | |  |
| 2223 | TOX21_PR_LUC_Followup_Antagonist | Follow-up assay | 110 | 73 | | 37 | |  | |  |  |  | |  |
| 2225 | TOX21_PR_LUC_Followup_Antagonist_viability | Follow-up assay | 169 | 0 | | 169 | |  | |  |  |  | |  |
| 2226 | TOX21_TR_LUC_GH3_Agonist_Followup | Follow-up assay | 33 | 5 | | 28 | |  | |  |  |  | |  |
| 2227 | TOX21_TR_LUC_GH3_Antagonist_Followup | Follow-up assay | 77 | 24 | | 53 | |  | |  |  |  | |  |
| 2253 | TOX21_TR_RXR_BLA_Agonist_Followup_ratio | Follow-up assay | 58 | 9 | | 49 | |  | |  |  |  | |  |
| 2254 | TOX21_TR_RXR_BLA_Agonist_Followup_viability | Follow-up assay | 63 | 2 | | 61 | |  | |  |  |  | |  |
| 2257 | TOX21_TR_RXR_BLA_Antagonist_Followup_ratio | Follow-up assay | 62 | 6 | | 56 | |  | |  |  |  | |  |
| 2258 | TOX21_TR_RXR_BLA_Antagonist_Followup_viability | Follow-up assay | 84 | 4 | | 80 | |  | |  |  |  | |  |
| 2230 | TOX21_TRA_COA_Agonist_Followup_ratio | Follow-up assay | 65 | 6 | | 59 | |  | |  |  |  | |  |
| 2233 | TOX21_TRA_COA_Antagonist_Followup_ratio | Follow-up assay | 81 | 0 | | 81 | |  | |  |  |  | |  |
| 2236 | TOX21_TRB_BLA_Agonist_Followup_ratio | Follow-up assay | 33 | 3 | | 30 | |  | |  |  |  | |  |
| 2237 | TOX21_TRB_BLA_Agonist_Followup_viability | Follow-up assay | 59 | 0 | | 59 | |  | |  |  |  | |  |
| 2240 | TOX21_TRB_BLA_Antagonist_Followup_ratio | Follow-up assay | 78 | 24 | | 54 | |  | |  |  |  | |  |
| 2241 | TOX21_TRB_BLA_Antagonist_Followup_viability | Follow-up assay | 65 | 0 | | 65 | |  | |  |  |  | |  |
| 2244 | TOX21_TRB_COA_Agonist_Followup_ratio | Follow-up assay | 66 | 9 | | 57 | |  | |  |  |  | |  |
| 2247 | TOX21_TRB_COA_Antagonist_Followup_ratio | Follow-up assay | 85 | 1 | | 84 | |  | |  |  |  | |  |
| 807 | TOX21_AhR_LUC_Agonist_viability | Viability | 5935 | 390 | | 5545 | |  | |  |  |  | |  |
| 1847 | TOX21_AP1_BLA_Agonist_viability | Viability | 5260 | 272 | | 4988 | |  | |  |  |  | |  |
| 763 | TOX21_AR_BLA_Antagonist_viability | Viability | 5141 | 487 | | 4654 | |  | |  |  |  | |  |
| 1823 | TOX21_AR_LUC_MDAKB2_Agonist_3uM_Nilutamide_viability | Viability | 6607 | 329 | | 6278 | |  | |  |  |  | |  |
| 1817 | TOX21_AR_LUC_MDAKB2_Antagonist_0.5nM_R1881_viability | Viability | 6751 | | 288 | | 6463 | |  |  |  |  | |  |
| 766 | TOX21_AR_LUC_MDAKB2_Antagonist_10nM_R1881_viability | Viability | 6296 | 387 | | 5909 | |  | |  |  |  | |  |
| 1185 | TOX21_ARE_BLA_agonist_viability | Viability | 5160 | 439 | | 4721 | |  | |  |  |  | |  |
| 768 | TOX21_Aromatase_LUC_Antagonist_viability | Viability | 5665 | 632 | | 5033 | |  | |  |  |  | |  |
| 2048 | TOX21_CAR_LUC_Agonist_viability | Viability | 5619 | 409 | | 5210 | |  | |  |  |  | |  |
| 2050 | TOX21_CAR_LUC_Antagonist_viability | Viability | 5828 | 457 | | 5371 | |  | |  |  |  | |  |
| 2369 | TOX21_CASP3_CHO_viability | Viability | 5129 | 910 | | 4219 | |  | |  |  |  | |  |
| 2367 | TOX21_CASP3_HEPG2_viability | Viability | 5901 | 322 | | 5579 | |  | |  |  |  | |  |
| 782 | TOX21_ELG1_LUC_Agonist_viability | Viability | 5569 | 228 | | 5341 | |  | |  |  |  | |  |
| 787 | TOX21_ERa_BLA_Antagonist_viability | Viability | 5708 | 189 | | 5519 | |  | |  |  |  | |  |
| 2212 | TOX21_ERa_LUC_VM7_Agonist_10nM_ICI182780_viability | Viability | 6230 | 357 | | 5873 | |  | |  |  |  | |  |
| 2054 | TOX21_ERa_LUC_VM7_Antagonist_0.1nM_E2_viability | Viability | 5627 | | 214 | | 5413 | |  |  |  |  | |  |
| 790 | TOX21_ERa_LUC_VM7_Antagonist_0.5nM_E2_viability | Viability | 5924 | | 291 | | 5633 | |  |  |  |  | |  |
| 2116 | TOX21_ERb_BLA_Agonist_viability | Viability | 4923 | 501 | | 4422 | |  | |  |  |  | |  |
| 2120 | TOX21_ERb_BLA_Antagonist_viability | Viability | 4480 | 468 | | 4012 | |  | |  |  |  | |  |
| 2059 | TOX21_ERR_LUC_viability | Viability | 5379 | 409 | | 4970 | |  | |  |  |  | |  |
| 1343 | TOX21_ESRE_BLA_Agonist_viability | Viability | 5239 | 255 | | 4984 | |  | |  |  |  | |  |
| 1188 | TOX21_FXR_BLA_agonist_viability | Viability | 5591 | 286 | | 5305 | |  | |  |  |  | |  |
| 1121 | TOX21_FXR_BLA_Antagonist_viability | Viability | 5181 | 348 | | 4833 | |  | |  |  |  | |  |
| 795 | TOX21_GR_BLA_Antagonist_viability | Viability | 5322 | 314 | | 5008 | |  | |  |  |  | |  |
| 2060 | TOX21_H2AX_HTRF_CHO_viability | Viability | 5967 | 363 | | 5604 | |  | |  |  |  | |  |
| 2062 | TOX21_HDAC_LUC_Antagonist_viability | Viability | 6333 | 75 | | 6258 | |  | |  |  |  | |  |
| 2066 | TOX21_HRE_BLA_Agonist_viability | Viability | 4676 | 470 | | 4206 | |  | |  |  |  | |  |
| 1186 | TOX21_HSE_BLA_agonist_viability | Viability | 5729 | 242 | | 5487 | |  | |  |  |  | |  |
| 799 | TOX21_MMP_viability | Viability | 5490 | 96 | | 5394 | |  | |  |  |  | |  |
| 1347 | TOX21_NFkB_BLA_agonist_viability | Viability | 3910 | 279 | | 3631 | |  | |  |  |  | |  |
| 1187 | TOX21_p53_BLA_p1_viability | Viability | 6271 | 265 | | 6006 | |  | |  |  |  | |  |
| 1318 | TOX21_p53_BLA_p2_viability | Viability | 3789 | 21 | | 3768 | |  | |  |  |  | |  |
| 1322 | TOX21_p53_BLA_p3_viability | Viability | 3520 | 13 | | 3507 | |  | |  |  |  | |  |
| 1326 | TOX21_p53_BLA_p4_viability | Viability | 3709 | 23 | | 3686 | |  | |  |  |  | |  |
| 1330 | TOX21_p53_BLA_p5_viability | Viability | 4547 | 13 | | 4534 | |  | |  |  |  | |  |
| 2072 | TOX21_PGC_ERR_LUC_viability | Viability | 5282 | 445 | | 4837 | |  | |  |  |  | |  |
| 1195 | TOX21_PPARd_BLA_Agonist_viability | Viability | 4631 | 578 | | 4053 | |  | |  |  |  | |  |
| 1126 | TOX21_PPARd_BLA_Antagonist_viability | Viability | 5093 | 414 | | 4679 | |  | |  |  |  | |  |
| 1128 | TOX21_PPARg_BLA_Antagonist_viability | Viability | 5247 | 317 | | 4930 | |  | |  |  |  | |  |
| 2124 | TOX21_PR_BLA_Agonist_viability | Viability | 4673 | 411 | | 4262 | |  | |  |  |  | |  |
| 2128 | TOX21_PR_BLA_Antagonist_viability | Viability | 4746 | 447 | | 4299 | |  | |  |  |  | |  |
| 2362 | TOX21_PXR_LUC_Agonist_viability | Viability | 5424 | 743 | | 4681 | |  | |  |  |  | |  |
| 1660 | TOX21_RAR_LUC_Agonist_viability | Viability | 6598 | 238 | | 6360 | |  | |  |  |  | |  |
| 1840 | TOX21_RAR_LUC_Antagonist_viability | Viability | 6929 | 169 | | 6760 | |  | |  |  |  | |  |
| 1662 | TOX21_RORg_LUC_CHO_Antagonist_viability | Viability | 5266 | 356 | | 4910 | |  | |  |  |  | |  |
| 1821 | TOX21_RXR_BLA_Agonist_viability | Viability | 4933 | 531 | | 4402 | |  | |  |  |  | |  |
| 2377 | TOX21_SBE_BLA_Agonist_viability | Viability | 5288 | 159 | | 5129 | |  | |  |  |  | |  |
| 2373 | TOX21_SBE_BLA_Antagonist_viability | Viability | 5246 | 161 | | 5085 | |  | |  |  |  | |  |
| 2107 | TOX21_SHH_3T3_GLI3_LUC_Agonist_viability | Viability | 5495 | 602 | | 4893 | |  | |  |  |  | |  |
| 2109 | TOX21_SHH_3T3_GLI3_LUC_Antagonist_viability | Viability | 5218 | 620 | | 4598 | |  | |  |  |  | |  |
| 805 | TOX21_TR_LUC_GH3_Antagonist_viability | Viability | 5174 | 1088 | | 4086 | |  | |  |  |  | |  |
| 1331 | TOX21_VDR_BLA_Agonist_viability | Viability | 4462 | 531 | | 3931 | |  | |  |  |  | |  |
| 1133 | TOX21_VDR_BLA_Antagonist_viability | Viability | 4821 | 355 | | 4466 | |  | |  |  |  | |  |
| 3190 | TOX21_AChE_Antagonist | Low positive | 5609 | 10 | | 5599 | |  | |  |  |  | |  |
| 2112 | TOX21_AChE_Fluor_Antagonist | Low positive | 7245 | 1 | | 7244 | |  | |  |  |  | |  |
| 3189 | TOX21_AChE_ms_Antagonist | Low positive | 5841 | 4 | | 5837 | |  | |  |  |  | |  |
| 764 | TOX21_AR_LUC_MDAKB2_Agonist | Low positive | 7185 | 191 | | 6994 | |  | |  |  |  | |  |
| 1822 | TOX21_AR_LUC_MDAKB2_Agonist_3uM_Nilutamide | Low positive | 6564 | 107 | | 6457 | | 246 | |  |  |  | |  |
| 2049 | TOX21_CAR_LUC_Antagonist | Low positive | 3906 | 438 | | 3468 | | 288 | |  |  |  | |  |
| 2368 | TOX21_CASP3_CHO | Low positive | 7472 | 40 | | 7432 | | 641 | |  |  |  | |  |
| 2366 | TOX21_CASP3_HEPG2 | Low positive | 7211 | 147 | | 7064 | | 167 | |  |  |  | |  |
| 781 | TOX21_ELG1_LUC_Agonist | Low positive | 7372 | 20 | | 7352 | | 123 | |  |  |  | |  |
| 2211 | TOX21_ERa_LUC_VM7_Agonist_10nM_ICI182780 | Low positive | 7102 | 89 | | 7013 | | 242 | |  |  |  | |  |
| 2115 | TOX21_ERb_BLA_Agonist_ratio | Low positive | 6309 | 50 | | 6259 | | 277 | |  |  |  | |  |
| 2055 | TOX21_ERR_LUC_Agonist | Low positive | 4771 | 129 | | 4642 | | 27 | |  |  |  | |  |
| 1342 | TOX21_ESRE_BLA_Agonist_ratio | Low positive | 6476 | 82 | | 6394 | | 101 | |  |  |  | |  |
| 1119 | TOX21_FXR_BLA_Agonist_ratio | Low positive | 6453 | 31 | | 6422 | | 109 | |  |  |  | |  |
| 1843 | TOX21_H2AX_HTRF_CHO_Agonist_ratio | Low positive | 6869 | 223 | | 6646 | | 179 | |  |  |  | |  |
| 2065 | TOX21_HRE_BLA_Agonist_ratio | Low positive | 7151 | 272 | | 6879 | | 349 | |  |  |  | |  |
| 1113 | TOX21_HSE_BLA_Agonist_ratio | Low positive | 6686 | 193 | | 6493 | | 150 | |  |  |  | |  |
| 2067 | TOX21_LUC_Biochem | Low positive | 6323 | 0 | | 6323 | |  | |  |  |  | |  |
| 796 | TOX21_MMP_fitc | Low positive | 6740 | 200 | | 6540 | | 30 | |  |  |  | |  |
| 798 | TOX21_MMP_rhodamine | Low positive | 4990 | 34 | | 4956 | | 6 | |  |  |  | |  |
| 1346 | TOX21_NFkB_BLA_agonist_ratio | Low positive | 6821 | 6 | | 6815 | | 92 | |  |  |  | |  |
| 1116 | TOX21_p53_BLA_p1_ratio | Low positive | 7236 | 294 | | 6942 | | 162 | |  |  |  | |  |
| 1317 | TOX21_p53_BLA_p2_ratio | Low positive | 4191 | 15 | | 4176 | | 8 | |  |  |  | |  |
| 1321 | TOX21_p53_BLA_p3_ratio | Low positive | 2972 | 12 | | 2960 | | 8 | |  |  |  | |  |
| 1325 | TOX21_p53_BLA_p4_ratio | Low positive | 3761 | 12 | | 3749 | | 6 | |  |  |  | |  |
| 1329 | TOX21_p53_BLA_p5_ratio | Low positive | 3983 | 11 | | 3972 | | 6 | |  |  |  | |  |
| 2068 | TOX21_PGC_ERR_LUC_Agonist | Low positive | 5684 | 96 | | 5588 | | 60 | |  |  |  | |  |
| 1124 | TOX21_PPARd_BLA_Agonist_ratio | Low positive | 5755 | 37 | | 5718 | | 188 | |  |  |  | |  |
| 1125 | TOX21_PPARd_BLA_Antagonist_ratio | Low positive | 5751 | 349 | | 5402 | | 242 | |  |  |  | |  |
| 802 | TOX21_PPARg_BLA_Agonist_ratio | Low positive | 6368 | 66 | | 6302 | |  | |  |  |  | |  |
| 2123 | TOX21_PR_BLA_Agonist_ratio | Low positive | 6716 | 132 | | 6584 | | 192 | |  |  |  | |  |
| 1659 | TOX21_RAR_LUC_Agonist | Low positive | 5524 | 12 | | 5512 | | 34 | |  |  |  | |  |
| 1820 | TOX21_RXR_BLA_Agonist_ratio | Low positive | 4154 | 130 | | 4024 | | 43 | |  |  |  | |  |
| 2376 | TOX21_SBE_BLA_Agonist_ratio | Low positive | 6582 | 0 | | 6582 | | 27 | |  |  |  | |  |
| 2106 | TOX21_SHH_3T3_GLI3_LUC_Agonist | Low positive | 6860 | 3 | | 6857 | | 311 | |  |  |  | |  |
| 803 | TOX21_TR_LUC_GH3_Agonist | Low positive | 7554 | 10 | | 7544 | |  | |  |  |  | |  |
| 2364 | TOX21_TRHR_HEK293_Agonist | Low positive | 6886 | 52 | | 6834 | |  | |  |  |  | |  |
| 2365 | TOX21_TRHR_HEK293_Antagonist | Low positive | 6195 | 81 | | 6114 | |  | |  |  |  | |  |
| 2043 | TOX21_TSHR_HTRF_Antagonist_ratio | Low positive | 6804 | 127 | | 6677 | |  | |  |  |  | |  |
| 2046 | TOX21_TSHR_wt_HTRF_Agonist_ratio | Low positive | 5737 | 23 | | 5714 | |  | |  |  |  | |  |
| 1131 | TOX21_VDR_BLA_Agonist_ratio | Low positive | 5343 | 15 | | 5328 | | 146 | |  |  |  | |  |
| 1132 | TOX21_VDR_BLA_Antagonist_ratio | Low positive | 5742 | 330 | | 5412 | | 267 | |  |  |  | |  |
| 2543 | TOX21_MSTI_p2_activity | No data | 0 | 0 | | 0 | |  | |  |  |  | |  |
| 2542 | TOX21_MSTI_p2_activity_n | No data | 0 | 0 | | 0 | |  | |  |  |  | |  |
| 3017 | TOX21_MSTI_p3_activity | No data | 0 | 0 | | 0 | |  | |  |  |  | |  |
| 3018 | TOX21_MSTI_p3_activity_n | No data | 0 | 0 | | 0 | |  | |  |  |  | |  |
| 2074 | TOX21_RT_HEK293_FLO_00hr_viability | Time-series data | 3311 | 2 | | 3309 | |  | |  |  |  | |  |
| 2075 | TOX21_RT_HEK293_FLO_08hr_viability | Time-series data | 2071 | 8 | | 2063 | |  | |  |  |  | |  |
| 2077 | TOX21_RT_HEK293_FLO_16hr_viability | Time-series data | 2066 | 11 | | 2055 | |  | |  |  |  | |  |
| 2078 | TOX21_RT_HEK293_FLO_24hr_viability | Time-series data | 1955 | 16 | | 1939 | |  | |  |  |  | |  |
| 2080 | TOX21_RT_HEK293_FLO_32hr_viability | Time-series data | 1946 | 23 | | 1923 | |  | |  |  |  | |  |
| 2082 | TOX21_RT_HEK293_FLO_40hr_viability | Time-series data | 1851 | 30 | | 1821 | |  | |  |  |  | |  |
| 2084 | TOX21_RT_HEK293_GLO_00hr_viability | Time-series data | 4023 | 0 | | 4023 | |  | |  |  |  | |  |
| 2086 | TOX21_RT_HEK293_GLO_08hr_viability | Time-series data | 2250 | 38 | | 2212 | |  | |  |  |  | |  |
| 2088 | TOX21_RT_HEK293_GLO_16hr_viability | Time-series data | 2080 | 42 | | 2038 | |  | |  |  |  | |  |
| 2089 | TOX21_RT_HEK293_GLO_24hr_viability | Time-series data | 2030 | 50 | | 1980 | |  | |  |  |  | |  |
| 2091 | TOX21_RT_HEK293_GLO_32hr_viability | Time-series data | 2080 | 51 | | 2029 | |  | |  |  |  | |  |
| 2093 | TOX21_RT_HEK293_GLO_40hr_viability | Time-series data | 2043 | 54 | | 1989 | |  | |  |  |  | |  |
| 2094 | TOX21_RT_HEPG2_FLO_00hr_viability | Time-series data | 3861 | 0 | | 3861 | |  | |  |  |  | |  |
| 2095 | TOX21_RT_HEPG2_FLO_08hr_viability | Time-series data | 3256 | 7 | | 3249 | |  | |  |  |  | |  |
| 2096 | TOX21_RT_HEPG2_FLO_16hr_viability | Time-series data | 2802 | 11 | | 2791 | |  | |  |  |  | |  |
| 2097 | TOX21_RT_HEPG2_FLO_24hr_viability | Time-series data | 2564 | 17 | | 2547 | |  | |  |  |  |  | |
| 2098 | TOX21_RT_HEPG2_FLO_32hr_viability | Time-series data | 2721 | 21 | | 2700 | |  | |  |  |  | |  |
| 2099 | TOX21_RT_HEPG2_FLO_40hr_viability | Time-series data | 2329 | 22 | | 2307 | |  | |  |  |  | |  |
| 2100 | TOX21_RT_HEPG2_GLO_00hr_viability | Time-series data | 3499 | 7 | | 3492 | |  | |  |  |  | |  |
| 2101 | TOX21_RT_HEPG2_GLO_08hr_viability | Time-series data | 2795 | 11 | | 2784 | |  | |  |  |  | |  |
| 2102 | TOX21_RT_HEPG2_GLO_16hr_viability | Time-series data | 2986 | 12 | | 2974 | |  | |  |  |  | |  |
| 2213 | TOX21_RT_HEPG2_GLO_24hr_viability | Time-series data | 3044 | 14 | | 3030 | |  | |  |  |  | |  |
| 2103 | TOX21_RT_HEPG2_GLO_32hr_viability | Time-series data | 3163 | 13 | | 3150 | |  | |  |  |  | |  |
| 2105 | TOX21_RT_HEPG2_GLO_40hr_viability | Time-series data | 3106 | 13 | | 3093 | |  | |  |  |  | |  |

**Supplementary Table 2: MACCS keys dictionary.** More information can be found at <https://www.mayachemtools.org/docs/modules/pdf/MACCSKeys.pdf>

| KEY | SMARTS | COUNT | DESCRIPTION |
| --- | --- | --- | --- |
| MACCS_1 | '?' | 0 | ISOTOPE |
| MACCS_1 | [#104] | 0 | 103 < ATOMIC NO. < 256 |
| MACCS_3 | '[#32,#33,#34,#50,#51,#52,#82,#83,#84]' | 0 | Group IVa,Va,VIa Rows 4-6 |
| MACCS_4 | '[Ac,Th,Pa,U,Np,Pu,Am,Cm,Bk,Cf,Es,Fm,Md,No,Lr]' | 0 | actinide |
| MACCS_5 | '[Sc,Ti,Y,Zr,Hf]' | 0 | Group IIIB,IVB (Sc...) |
| MACCS_6 | '[La,Ce,Pr,Nd,Pm,Sm,Eu,Gd,Tb,Dy,Ho,Er,Tm,Yb,Lu]' | 0 | Lanthanide |
| MACCS_7 | '[V,Cr,Mn,Nb,Mo,Tc,Ta,W,Re]' | 0 | Group VB,VIB,VIIB |
| MACCS_8 | '[!#6;!#1]1~*~*~*~1' | 0 | QAAA@1 |
| MACCS_9 | '[Fe,Co,Ni,Ru,Rh,Pd,Os,Ir,Pt]' | 0 | Group VIII (Fe...) |
| MACCS_10 | '[Be,Mg,Ca,Sr,Ba,Ra]' | 0 | Group IIa (Alkaline earth) |
| MACCS_11 | '*1~*~*~*~1' | 0 | 4M Ring |
| MACCS_12 | '[Cu,Zn,Ag,Cd,Au,Hg]' | 0 | Group IB,IIB (Cu..) |
| MACCS_13 | '[#8]~[#7](~[#6])~[#6]' | 0 | ON(C)C |
| MACCS_14 | '[#16]-[#16]' | 0 | S-S |
| MACCS_15 | '[#8]~[#6](~[#8])~[#8]' | 0 | OC(O)O |
| MACCS_16 | '[!#6;!#1]1~*~*~1' | 0 | QAA@1 |
| MACCS_17 | '[#6]#[#6]' | 0 | CTC |
| MACCS_18 | '[#5,#13,#31,#49,#81]' | 0 | Group IIIA (B...) |
| MACCS_19 | '*1~*~*~*~*~*~*~1' | 0 | 7M Ring |
| MACCS_30 | '[#6]~[!#6;!#1](~[#6])(~[#6])~*' | 0 | CQ(C)(C)A |
| MACCS_31 | '[!#6;!#1]~[F,Cl,Br,I]' | 0 | QX |
| MACCS_32 | '[#6]~[#16]~[#7]' | 0 | CSN |
| MACCS_33 | '[#7]~[#16]' | 0 | NS |
| MACCS_34 | '[CH2]=*' | 0 | CH2=A |
| MACCS_35 | '[Li,Na,K,Rb,Cs,Fr]' | 0 | Group IA (Alkali Metal) |
| MACCS_36 | '[#16R]' | 0 | S Heterocycle |
| MACCS_37 | '[#7]~[#6](~[#8])~[#7]' | 0 | NC(O)N |
| MACCS_38 | '[#7]~[#6](~[#6])~[#7]' | 0 | NC(C)N |
| MACCS_39 | '[#8]~[#16](~[#8])~[#8]' | 0 | OS(O)O |
| MACCS_40 | '[#16]-[#8]' | 0 | S-O |
| MACCS_41 | '[#6]#[#7]' | 0 | CTN |
| MACCS_42 | 'F' | 0 | F |
| MACCS_43 | '[!#6;!#1;!H0]~*~[!#6;!#1;!H0]' | 0 | QHAQH |
| MACCS_44 | '[!#1;!#6;!#7;!#8;!#9;!#14;!#15;!#16;!#17;!#35;!#53]' | 0 | OTHER |
| MACCS_45 | '[#6]=[#6]~[#7]' | 0 | C=CN |
| MACCS_46 | 'Br' | 0 | BR |
| MACCS_47 | '[#16]~*~[#7]' | 0 | SAN |
| MACCS_48 | '[#8]~[!#6;!#1](~[#8])(~[#8])' | 0 | OQ(O)O |
| MACCS_49 | '[!+0]' | 0 | CHARGE |
| MACCS_50 | '[#6]=[#6](~[#6])~[#6]' | 0 | C=C(C)C |
| MACCS_51 | '[#6]~[#16]~[#8]' | 0 | CSO |
| MACCS_52 | '[#7]~[#7]' | 0 | NN |
| MACCS_53 | '[!#6;!#1;!H0]~*~*~*~[!#6;!#1;!H0]' | 0 | QHAAAQH |
| MACCS_54 | '[!#6;!#1;!H0]~*~*~[!#6;!#1;!H0]' | 0 | QHAAQH |
| MACCS_55 | '[#8]~[#16]~[#8]' | 0 | OSO |
| MACCS_56 | '[#8]~[#7](~[#8])~[#6]' | 0 | ON(O)C |
| MACCS_57 | '[#8R]' | 0 | O Heterocycle |
| MACCS_58 | '[!#6;!#1]~[#16]~[!#6;!#1]' | 0 | QSQ |
| MACCS_59 | '[#16]!:*:*' | 0 | Snot%A%A |
| MACCS_60 | '[#16]=[#8]' | 0 | S=O |
| MACCS_61 | '*~[#16](~*)~*' | 0 | AS(A)A |
| MACCS_62 | '*@*!@*@*' | 0 | A$!A$A |
| MACCS_63 | '[#7]=[#8]' | 0 | N=O |
| MACCS_64 | '*@*!@[#16]' | 0 | A$A!S |
| MACCS_65 | 'c:n' | 0 | C%N |
| MACCS_66 | '[#6]~[#6](~[#6])(~[#6])~*' | 0 | CC(C)(C)A |
| MACCS_67 | '[!#6;!#1]~[#16]' | 0 | QS |
| MACCS_68 | '[!#6;!#1;!H0]~[!#6;!#1;!H0]' | 0 | QHQH (&...) SPEC Incomplete |
| MACCS_69 | '[!#6;!#1]~[!#6;!#1;!H0]' | 0 | QQH |
| MACCS_70 | '[!#6;!#1]~[#7]~[!#6;!#1]' | 0 | QNQ |
| MACCS_71 | '[#7]~[#8]' | 0 | NO |
| MACCS_72 | '[#8]~*~*~[#8]' | 0 | OAAO |
| MACCS_73 | '[#16]=*' | 0 | S=A |
| MACCS_74 | '[CH3]~*~[CH3]' | 0 | CH3ACH3 |
| MACCS_75 | '*!@[#7]@*' | 0 | A!N$A |
| MACCS_76 | '[#6]=[#6](~*)~*' | 0 | C=C(A)A |
| MACCS_77 | '[#7]~*~[#7]' | 0 | NAN |
| MACCS_78 | '[#6]=[#7]' | 0 | C=N |
| MACCS_79 | '[#7]~*~*~[#7]' | 0 | NAAN |
| MACCS_80 | '[#7]~*~*~*~[#7]' | 0 | NAAAN |
| MACCS_81 | '[#16]~*(~*)~*' | 0 | SA(A)A |
| MACCS_82 | '*~[CH2]~[!#6;!#1;!H0]' | 0 | ACH2QH |
| MACCS_83 | '[!#6;!#1]1~*~*~*~*~1' | 0 | QAAAA@1 |
| MACCS_84 | '[NH2]' | 0 | NH2 |
| MACCS_85 | '[#6]~[#7](~[#6])~[#6]' | 0 | CN(C)C |
| MACCS_86 | '[C;H2,H3][!#6;!#1][C;H2,H3]' | 0 | CH2QCH2 |
| MACCS_87 | '[F,Cl,Br,I]!@*@*' | 0 | X!A$A |
| MACCS_88 | '[#16]' | 0 | S |
| MACCS_89 | '[#8]~*~*~*~[#8]' | 0 | OAAAO |
| MACCS_90 | '[$([!#6;!#1;!H0]~*~*~[CH2]~*),$([!#6;!#1;!H0;R]1@[R]@[R]@[CH2;R]1),$([!#6;!#1;!H0]~[R]1@[R]@[CH2;R]1)]' | 0 | QHAACH2A |
| MACCS_91 | '[$([!#6;!#1;!H0]~*~*~*~[CH2]~*),$([!#6;!#1;!H0;R]1@[R]@[R]@[R]@[CH2;R]1),$([!#6;!#1;!H0]~[R]1@[R]@[R]@[CH2;R]1),$([!#6;!#1;!H0]~*~[R]1@[R]@[CH2;R]1)]' | 0 | QHAAACH2A |
| MACCS_92 | '[#8]~[#6](~[#7])~[#6]' | 0 | OC(N)C |
| MACCS_93 | '[!#6;!#1]~[CH3]' | 0 | QCH3 |
| MACCS_94 | '[!#6;!#1]~[#7]' | 0 | QN |
| MACCS_95 | '[#7]~*~*~[#8]' | 0 | NAAO |
| MACCS_96 | '*1~*~*~*~*~1' | 0 | 5 M ring |
| MACCS_97 | '[#7]~*~*~*~[#8]' | 0 | NAAAO |
| MACCS_98 | '[!#6;!#1]1~*~*~*~*~*~1' | 0 | QAAAAA@1 |
| MACCS_99 | '[#6]=[#6]' | 0 | C=C |
| MACCS_100 | '*~[CH2]~[#7]' | 0 | ACH2N |
| MACCS_101 | '[$([R]@1@[R]@[R]@[R]@[R]@[R]@[R]@[R]1),$([R]@1@[R]@[R]@[R]@[R]@[R]@[R]@[R]@[R]1),$([R]@1@[R]@[R]@[R]@[R]@[R]@[R]@[R]@[R]@[R]1),$([R]@1@[R]@[R]@[R]@[R]@[R]@[R]@[R]@[R]@[R]@[R]1),$([R]@1@[R]@[R]@[R]@[R]@[R]@[R]@[R]@[R]@[R]@[R]@[R]1),$([R]@1@[R]@[R]@[R]@[R]@[R]@[R]@[R]@[R]@[R]@[R]@[R]@[R]1),$([R]@1@[R]@[R]@[R]@[R]@[R]@[R]@[R]@[R]@[R]@[R]@[R]@[R]@[R]1)]' | 0 | 8M Ring or larger. This only handles up to ring sizes of 14 |
| MACCS_102 | '[!#6;!#1]~[#8]' | 0 | QO |
| MACCS_103 | 'Cl' | 0 | CL |
| MACCS_104 | '[!#6;!#1;!H0]~*~[CH2]~*' | 0 | QHACH2A |
| MACCS_105 | '*@*(@*)@*' | 0 | A$A($A)$A |
| MACCS_106 | '[!#6;!#1]~*(~[!#6;!#1])~[!#6;!#1]' | 0 | QA(Q)Q |
| MACCS_107 | '[F,Cl,Br,I]~*(~*)~*' | 0 | XA(A)A |
| MACCS_108 | '[CH3]~*~*~*~[CH2]~*' | 0 | CH3AAACH2A |
| MACCS_109 | '*~[CH2]~[#8]' | 0 | ACH2O |
| MACCS_110 | '[#7]~[#6]~[#8]' | 0 | NCO |
| MACCS_111 | '[#7]~*~[CH2]~*' | 0 | NACH2A |
| MACCS_112 | '*~*(~*)(~*)~*' | 0 | AA(A)(A)A |
| MACCS_113 | '[#8]!:*:*' | 0 | Onot%A%A |
| MACCS_114 | '[CH3]~[CH2]~*' | 0 | CH3CH2A |
| MACCS_115 | '[CH3]~*~[CH2]~*' | 0 | CH3ACH2A |
| MACCS_116 | '[$([CH3]~*~*~[CH2]~*),$([CH3]~*1~*~[CH2]1)]' | 0 | CH3AACH2A |
| MACCS_117 | '[#7]~*~[#8]' | 0 | NAO |
| MACCS_118 | '[$(*~[CH2]~[CH2]~*),$(*1~[CH2]~[CH2]1)]' | 1 | ACH2CH2A > 1 |
| MACCS_119 | '[#7]=*' | 0 | N=A |
| MACCS_120 | '[!#6;R]' | 1 | Heterocyclic atom > 1 (&...) Spec Incomplete |
| MACCS_121 | '[#7;R]' | 0 | N Heterocycle |
| MACCS_122 | '*~[#7](~*)~*' | 0 | AN(A)A |
| MACCS_123 | '[#8]~[#6]~[#8]' | 0 | OCO |
| MACCS_124 | '[!#6;!#1]~[!#6;!#1]' | 0 | QQ |
| MACCS_125 | '?' | 0 | Aromatic Ring > 1 |
| MACCS_126 | '*!@[#8]!@*' | 0 | A!O!A |
| MACCS_127 | '*@*!@[#8]' | 1 | A$A!O > 1 (&...) Spec Incomplete |
| MACCS_128 | '[$(*~[CH2]~*~*~*~[CH2]~*),$([R]1@[CH2;R]@[R]@[R]@[R]@[CH2;R]1),$(*~[CH2]~[R]1@[R]@[R]@[CH2;R]1),$(*~[CH2]~*~[R]1@[R]@[CH2;R]1)]' | 0 | ACH2AAACH2A |
| MACCS_129 | '[$(*~[CH2]~*~*~[CH2]~*),$([R]1@[CH2]@[R]@[R]@[CH2;R]1),$(*~[CH2]~[R]1@[R]@[CH2;R]1)]' | 0 | ACH2AACH2A |
| MACCS_130 | '[!#6;!#1]~[!#6;!#1]' | 1 | QQ > 1 (&...) Spec Incomplete |
| MACCS_131 | '[!#6;!#1;!H0]' | 1 | QH > 1 |
| MACCS_132 | '[#8]~*~[CH2]~*' | 0 | OACH2A |
| MACCS_133 | '*@*!@[#7]' | 0 | A$A!N |
| MACCS_134 | '[F,Cl,Br,I]' | 0 | X (HALOGEN) |
| MACCS_135 | '[#7]!:*:*' | 0 | Nnot%A%A |
| MACCS_136 | '[#8]=*' | 1 | O=A>1 |
| MACCS_137 | '[!C;!c;R]' | 0 | Heterocycle |
| MACCS_138 | '[!#6;!#1]~[CH2]~*' | 1 | QCH2A>1 (&...) Spec Incomplete |
| MACCS_139 | '[O;!H0]' | 0 | OH |
| MACCS_140 | '[#8]' | 3 | O > 3 (&...) Spec Incomplete |
| MACCS_141 | '[CH3]' | 2 | CH3 > 2 (&...) Spec Incomplete |
| MACCS_142 | '[#7]' | 1 | N > 1 |
| MACCS_143 | '*@*!@[#8]' | 0 | A$A!O |
| MACCS_144 | '*!:*:*!:*' | 0 | Anot%A%Anot%A |
| MACCS_145 | '*1~*~*~*~*~*~1' | 1 | 6M ring > 1 |
| MACCS_146 | '[#8]' | 2 | O > 2 |
| MACCS_147 | '[$(*~[CH2]~[CH2]~*),$([R]1@[CH2;R]@[CH2;R]1)]' | 0 | ACH2CH2A |
| MACCS_148 | '*~[!#6;!#1](~*)~*' | 0 | AQ(A)A |
| MACCS_149 | '[C;H3,H4]' | 1 | CH3 > 1 |
| MACCS_150 | '*!@*@*!@*' | 0 | A!A$A!A |
| MACCS_151 | '[#7;!H0]' | 0 | NH |
| MACCS_152 | '[#8]~[#6](~[#6])~[#6]' | 0 | OC(C)C |
| MACCS_153 | '[!#6;!#1]~[CH2]~*' | 0 | QCH2A |
| MACCS_154 | '[#6]=[#8]' | 0 | C=O |
| MACCS_155 | '*!@[CH2]!@*' | 0 | A!CH2!A |
| MACCS_156 | '[#7]~*(~*)~*' | 0 | NA(A)A |
| MACCS_157 | '[#6]-[#8]' | 0 | C-O |
| MACCS_158 | '[#6]-[#7]' | 0 | C-N |
| MACCS_159 | '[#8]' | 1 | O>1 |
| MACCS_160 | '[C;H3,H4]' | 0 | CH3 |
| MACCS_161 | '[#7]' | 0 | N |
| MACCS_162 | 'a' | 0 | Aromatic |
| MACCS_163 | '*1~*~*~*~*~*~1' | 0 | 6M Ring |
| MACCS_164 | '[#8]' | 0 | O |
| MACCS_165 | '[R]' | 0 | Ring |
| MACCS_166 | '?' | 0 | Fragments FIX: this can't be done in SMARTS |
